# Engineering functional ventral midbrain dopaminergic neurons in human organoids through WNT modulation and bioreactor culture

**DOI:** 10.1101/2025.07.29.667404

**Authors:** Hariam Raji, Federico Bertoli, Maria Jose Perez, Alicia Lam, Laura Volpicelli-Daley, Michela Deleidi

## Abstract

Human midbrain organoids (hMOs) derived from induced pluripotent stem cells provide a powerful system to model disorders involving dopamine (DA) dysfunction, including Parkinson’s disease (PD) and neuropsychiatric conditions. However, current differentiation protocols still fall short in recapitulating early specification, substantia nigra pars compacta (SNpc)-like identity, and the functional maturation of vulnerable DA neurons. Here, we established a differentiation strategy that combines tri-phasic WNT modulation with dynamic bioreactor culture to generate hMOs enriched in SNpc-like DA neurons. This approach significantly increases the yield of TH⁺/GIRK2⁺ and TH⁺/ALDH1A1⁺ DA neurons and promotes enhanced synaptic maturation, robust electrophysiological activity, and elevated DA release. Single-cell transcriptomics revealed that this strategy drives the emergence of *SOX6*^+^/*GIRK2*^+^ SNpc-like neurons, accompanied by upregulation of synaptic, metabolic, and maturation programs, alongside reduced cell stress and apoptotic signaling. Importantly, hMOs demonstrated vulnerability upon exposure to α-synuclein preformed fibrils, resulting in aggregate formation and DA neuron degeneration, supporting their use as a human model of PD-relevant pathology. Overall, this system provides a scalable and physiologically relevant approach to investigate molecular mechanisms underlying neurodegeneration and DA-related disorders.

## Introduction

The development of three-dimensional (3D) human brain organoids has transformed the landscape of neurological disease modeling by offering unprecedented opportunities to study complex tissue architecture, cell interactions, and developmental processes *in vitro*^1–4^. Organoids derived from human induced pluripotent stem cells (iPSCs) can self-organize into structures that recapitulate key aspects of brain development, including spatial patterning, region-specific differentiation, and organ-like cytoarchitectural organization^5^. These systems have been instrumental in modeling neurodevelopmental disorders and are increasingly recognized as valuable tools for studying neurodegenerative diseases. They also offer promising opportunities for personalized medicine, including patient-specific drug testing and cell-based therapeutic strategies^6^.

Among brain organoid types, human midbrain organoids (hMOs) have emerged as particularly promising for studying diseases that affect dopamine (DA) neurons, such as Parkinson’s disease (PD), owing to their ability to mimic the early development and regional identity of the human ventral midbrain (VM), which harbors disease-relevant DA subtypes^7, 8^ . Within the VM, DA neurons are organized into two principal groups: A9 neurons located in the SNpc and A10 neurons in the medial and ventral tegmental area (VTA)^9, 10^. In the context of PD, A9 DA neurons are consistently more susceptible to degeneration, while A10 DA neurons are relatively spared^11, 12^. Vulnerable SNpc DA neurons arise from midbrain floor plate progenitors (FPP) during early embryogenesis, with their specification and maturation guided by morphogen gradients, including SHH, FGF8, and WNT signaling^13, 14^. Among these, canonical WNT/β-catenin signaling plays a central role by regulating key transcription factors such as EN1, LMX1A/B, and OTX2 ^15–19^. These developmental cues also shape DA subtype identity along the VTA-SNpc axis, as reflected by region-specific markers including EN1/2, ALDH1A1, GIRK2, SOX6, and EBF1^20, 21^.

Despite these advances, generating hMOs enriched in mature and functionally relevant midbrain DA (mDA) neurons remains a major challenge. Current differentiation protocols often lack the precision required to recapitulate the molecular and functional features of selectively vulnerable DA neuron subtypes^22, 23^. In addition, conventional orbital shaking systems are limited by insufficient oxygen and nutrient diffusion^24^, particularly as organoids increase in size and complexity. These constraints hinder the maturation of metabolically demanding neuronal populations, such as SNpc DA neurons, which are especially vulnerable in PD^11, 25, 26^. To address these limitations, we developed a differentiation strategy that combines temporally controlled WNT signaling with dynamic spinning bioreactor culture. This tri-phasic WNT modulation approach, implemented in a scalable bioreactor system, is designed to more closely mimic key *in vivo* developmental transitions governing DA neuron specification and maturation. Specifically, an initial phase of WNT activation promotes expansion of midbrain FPP, followed by a precisely timed inhibition of WNT signaling to drive DA precursor specification and terminal differentiation. In parallel, spinning bioreactors enhance oxygen and nutrient delivery throughout the 3D structure, supporting uniform cell survival, improved maturation, and increased cytoarchitectural complexity of the developing neuronal niche.

Collectively, this approach enables the generation of a more physiologically relevant organoid model, providing a platform to investigate the mechanisms underlying selective SNpc DA neuron vulnerability and helping to address a critical gap in the field.

## Results

### Tri-phasic WNT modulation promotes midbrain patterning and early DA specification in human midbrain organoids

To promote the specification of human iPSC-derived DA neurons toward a SNpc lineage in hMOs, we precisely fine-tuned WNT signaling dynamics, a pathway critical for early midbrain patterning, floor plate identity, and DA lineage commitment^15, 17, 27, 28^. We developed a tri-phasic WNT modulation strategy (WntTriP), involving an initial bi-phasic WNT activation followed by inhibition with IWP-2, and implemented it using orbital shakers (OS), referred to as the ‘WntTriP-OS’ protocol. This was compared with our previously published differentiation protocol^7^, which we used as a baseline to refine and enrich VM identity, here referred to as the Standard-OS condition **(Figure 1A-E)**. Immunofluorescence analysis at day *in vitro* (DIV) 20 confirmed neural induction and midbrain specification, with expression of SOX2, FOXA2, OTX2, EN1, and LMX1A (**Figure 1F**). Importantly, WntTriP-OS organoids exhibited increased expression of tyrosine hydroxylase (TH) compared to Standard-OS (**Figure 1G,H**), indicating enhanced early DA neuron production. These findings were reproducible across additional iPSC lines (**Supplementary Figure 1A,B**).

**Figure 1.**
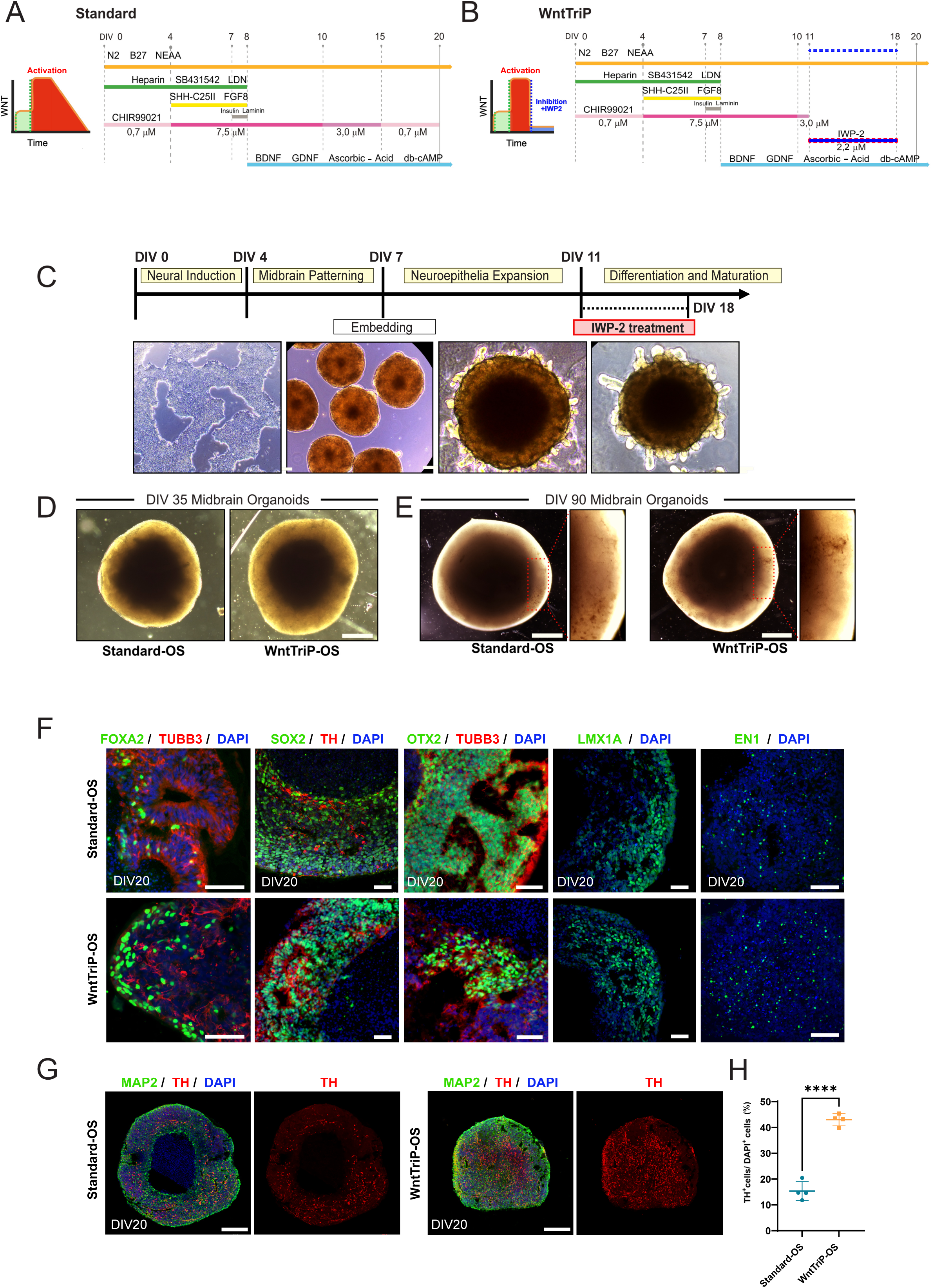
Generation and characterization of midbrain organoids through tri-phasic modulation of WNT signaling. **(A, B)** Schematic diagrams depicting the patterning factors and differentiation timelines for the Standard and tri-phasic WNT (WntTriP) protocols used to generate human midbrain organoids (hMOs). **(C)** Brightfield images illustrating Matrigel embedding and IWP-2 treatment stages during WntTriP-OS organoid differentiation. Scale bars: 250 µm. **(D)** Representative brightfield images of Standard-OS and WntTriP-OS hMOs at DIV 35. Scale bars: 500 µm. **(E)** Representative brightfield images of Standard-OS and WntTriP-OS hMOs at DIV 90. Adjacent panels show higher magnification images. Scale bars: 1000 µm. **(F)** Representative immunofluorescence images of DIV 20 hMOs showing expression of midbrain floor plate markers FOXA2 (green), OTX2 (green), midbrain DA progenitor markers LMX1A (green) and EN1 (green), the DA marker tyrosine hydroxylase (TH, red), the pan-neuronal marker TUBB3 (red) and the neuronal progenitor marker SOX2 (green). Cell nuclei were counterstained with DAPI (blue). Scale bars: 20 µm. **(G)** Representative immunofluorescence images of hMOs at DIV 20, showing expression of MAP2 (green) and TH (red). Cell nuclei were counterstained with DAPI (blue). Scale bars: 200 µm. **(H)** Quantification of TH^+^ neurons expressed as the percentage of total DAPI^+^ cells in hMO sections at DIV 20. MeanLJ±LJSD; unpaired two-tailed t-test; ****p < 0.0001; n=4 organoids per condition.

### Single-cell transcriptomic analysis reveals enhanced ventral midbrain patterning and SNpc-like DA specification following tri-phasic WNT modulation

To assess the developmental context of the WNT modulation, we performed single-cell RNA sequencing (scRNA-seq) at DIV25. Label transfer using the Braun et al. human fetal brain atlas demonstrated predominant mapping to midbrain populations, with minor contributions from diencephalic and medullary identities (**Figure 2A, Supplementary Datasheet 1**). Developmental projection placed most cells in PCW14-like states, corresponding to early second-trimester midbrain development (**Supplementary Figure 2A**). Unsupervised clustering identified diverse neurodevelopmental populations, including radial glia, neural progenitors, FPP, DA neuroblasts, differentiating and mature DA neurons, VM GABAergic neurons, and a distinct *SOX6*⁺/*EBF1*⁺ VM DA neuron cluster (**Figure 2B-D, Supplementary Datasheet 1**). Marker analysis confirmed appropriate lineage progression, with *SOX2* enriched in progenitors, *DCX* in neuronal populations, and VM markers (*EN1*, *EBF1*, *NR4A2*, *CNTN5*, *PCDH7*, *SOX6*) along DA trajectories. Cluster composition was consistent across replicates (**Supplementary Figure 2B**).

**Figure 2.**
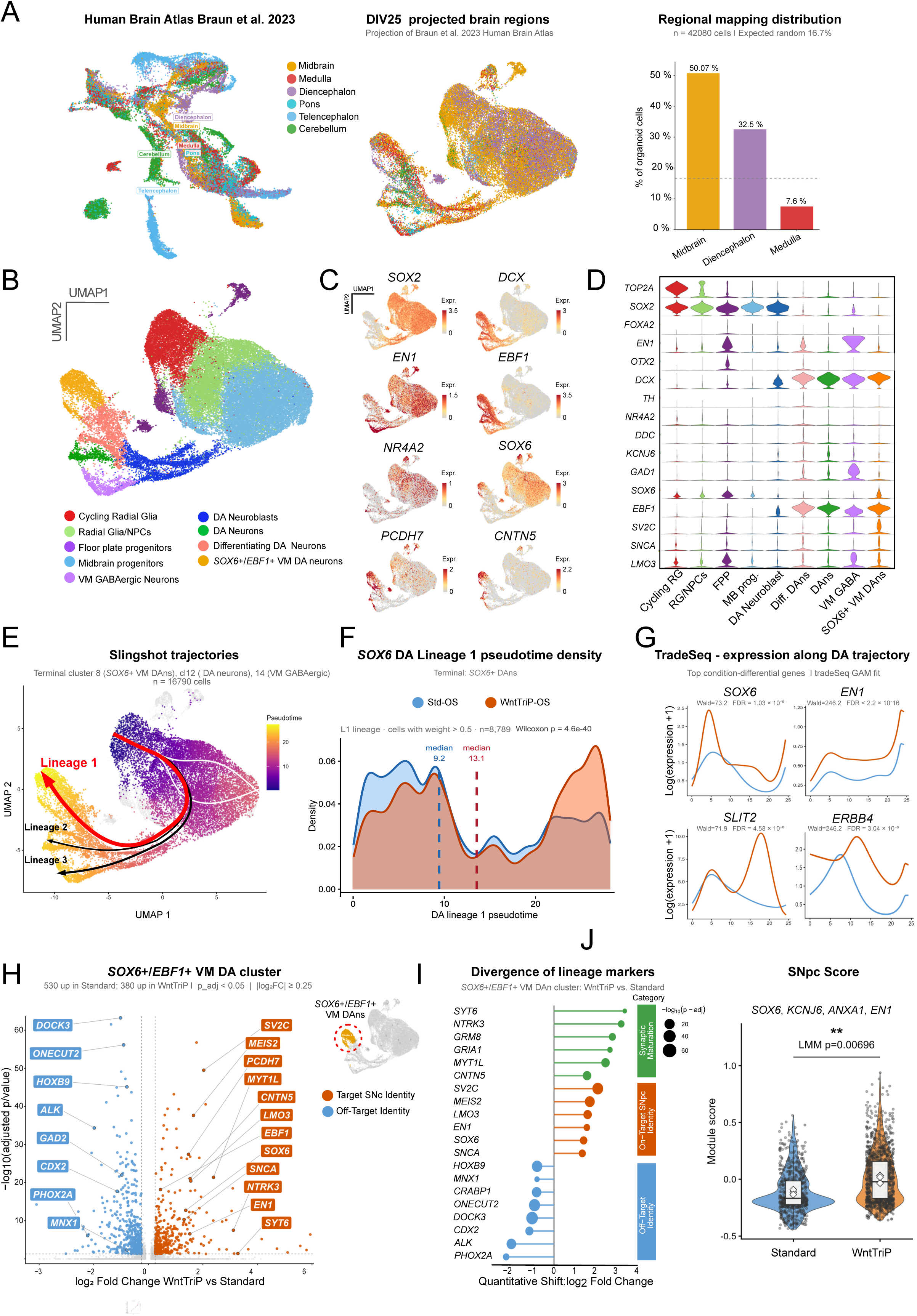
Single-cell transcriptomic analysis of DIV25 organoids reveals enhanced ventral midbrain and SNpc-like dopaminergic (DA) specification following tri-phasic WNT modulation. **(A)** Label transfer projection of the Braun et al. (2023) human brain atlas^85^ onto DIV25 hMO cells. Left: reference brain regions. Middle: projected regional identities on the DIV25 UMAP. Right: predominant assignment to midbrain identity. **(B)** UMAP visualization of the integrated DIV25 single-cell dataset from Standard and WntTriP organoids, colored by cluster annotation, including radial glia (RG), RGs/NPCs, floor plate progenitors (FPP), midbrain progenitors, DA neuroblasts, differentiating DA neurons, DA neurons, *SOX6*⁺/*EBF1*⁺ VM DA neurons, and VM GABAergic neurons. **(C)** Feature plots showing expression of representative lineage and DA markers (*SOX2*, *DCX*, *EN1*, *EBF1*, *NR4A2*, *PCDH7*, *CNTN5*, *SOX6*) across the UMAP. **(D)** Violin plots illustrating the distribution of selected marker genes across annotated clusters. **(E)** Slingshot trajectory inference across DIV25 DA lineages. Arrows indicate inferred developmental paths terminating in immature DA neurons, DA neurons, and the lineage of interest, *SOX6*⁺ DA neurons (Lineage 1, red). **(F)** Pseudotime density plot for *SOX6*⁺ lineage 1, showing a rightward shift in WntTriP organoids, consistent with accelerated progression toward this lineage. **(G)** TradeSeq (fitGAM) smoothed gene expression dynamics along the DA trajectory for representative condition-dependent genes (*SOX6*, *EN1*, *SLIT2*, *ERBB4*). **(H)** Differential gene expression analysis of the *SOX6*⁺/*EBF1*⁺ DA neuron cluster comparing WntTriP and Standard organoids at DIV25, showing enrichment of substantia nigra-associated genes in WntTriP and off-target lineage markers in Standard organoids. **(I)** Quantitative comparison of lineage marker divergence within the *SOX6*⁺/*EBF1*⁺ DA cluster, highlighting genes associated with synaptic maturation, SNpc identity, and off-target lineages. **(J)** Violin plot of SNpc module scores (*SOX6*, *KCNJ6*, *ANXA1*, *EN1*) in *SOX6*⁺/*EBF1*⁺ DA neurons, showing significantly higher scores in WntTriP organoids. N = 3 hMOs; LMM, **p = 0.00696.

Next, we assessed whether tri-phasic WNT modulation influenced cell-state progression and endpoint specification within the DA differentiation trajectory. Trajectory inference (Slingshot) revealed two DA differentiation lineages, with one terminating in the *SOX6*⁺/*EBF1*⁺ VM DA cluster (**Figure 2E**). WntTriP-OS organoids showed increased enrichment at late pseudotime along this lineage, indicating enhanced progression toward this endpoint (**Figure 2F**). Consistently, tradeSeq analysis along the DA lineage identified increased expression of neuronal maturation and SNpc-associated genes, including *SOX6*, *EN1*, *SLIT2*, and *ERBB4*, under WntTriP conditions (**Figure 2G**).

Focusing on the *SOX6*⁺/*EBF1*⁺ cluster, differential expression analysis revealed enrichment of a VM/SNpc-like transcriptional program in WntTriP organoids, including *SOX6, EBF1, EN1, LMO3*, and *SV2C* ^29–32^, alongside genes linked to synaptic maturation (*CNTN5, SYT6, SNCA, NTRK3*). In contrast, Standard-OS organoids showed higher expression of posterior or non-midbrain markers (e.g., *CDX2, HOXB9, ONECUT2, PHOX2A, MNX1, ALK*), consistent with alternative lineage identities ^33–35^ (**Figure 2H,I; Supplementary Datasheet 1**). A strict SNpc gene module score (*SOX6, KCNJ6, ANXA1, EN1*) was significantly increased in the *SOX6*⁺/*EBF1*⁺ cluster under WntTriP conditions (**Figure 2J**). Together, these results indicate that WntTriP enhances both DA neuron production and specification toward an SNpc-like identity.

To examine the signaling environment underlying this specification, we performed cell-cell communication analysis using CellChat across annotated populations at DIV25. WntTriP organoids exhibited reduced non-canonical WNT signaling and a reorganization of FGF signaling centered on the *SOX6*⁺/*EBF1*⁺ VM DA cluster **(Supplementary Figure 2C)**. Ligand–receptor analysis revealed increased interactions in axon guidance (*FLRT2*-*UNC5D*, *EFNB2*/3-*EPHB1*), synaptic adhesion (*CNTN1*-*NRCAM*, *NCAM1*-*NCAM2*), and VM patterning pathways (*BMP5-BMPR1A/BMPR2*, *DLL3-NOTCH2*) **(Supplementary Figure 2D)**. Pathway directionality analysis further revealed strong incoming SLIT signaling to the *SOX6*⁺/*EBF1*⁺ cluster, consistent with SLIT-ROBO axonal guidance cues shaping the emerging SNpc-like environment^36, 37^. In parallel, ADGRL signaling, a synapse-organizing G protein-coupled receptor^38^, was highly bidirectional, suggesting active synaptogenic interactions via ADGRL-FLRT pathways^39, 40^ **(Supplementary Figure 2E)**. Collectively, these findings demonstrate that tri-phasic WNT modulation not only enhances early DA neuron production but also promotes a transcriptional and signaling environment, including synaptogenic programs, which favor SNpc-like specification and early maturation.

### Dynamic bioreactor culture enhances tissue viability and dopaminergic neuron survival

To assess the impact of WntTriP modulation on DA neuron maturation, we first examined VMAT2, a marker of functional DA neurons involved in vesicular DA transport ^41^. At DIV65, WntTriP-OS organoids showed increased VMAT2 expression compared to Standard-OS; however, VMAT2 only partially co-localized with TH and did not significantly differ between conditions at the single-cell level (**Figure 3A,B**). We next evaluated GIRK2, a marker of SNpc-like DA neurons ^42^. GIRK2 expression was detected in TH⁺ neurons in WntTriP-OS organoids but was absent in Standard-OS (**Figure 3C**). Analysis of ALDH1A1, associated with mature SNpc DA neurons ^43, 44^, revealed low expression in both conditions at DIV40, but a marked increase in WntTriP-OS organoids by DIV100 (**Figure 3D**). Notably, this increase coincided with a substantial loss of TH⁺ neurons, suggesting impaired long-term maintenance of the DA phenotype. Given that ALDH1A1 is not DA-specific ^45^, its late expression likely does not reflect functional DA neuron preservation. Together, these findings indicate that WntTriP accelerates DA differentiation and promotes SNpc-like identity, but compromises long-term stability under orbital shaking conditions.

**Figure 3.**
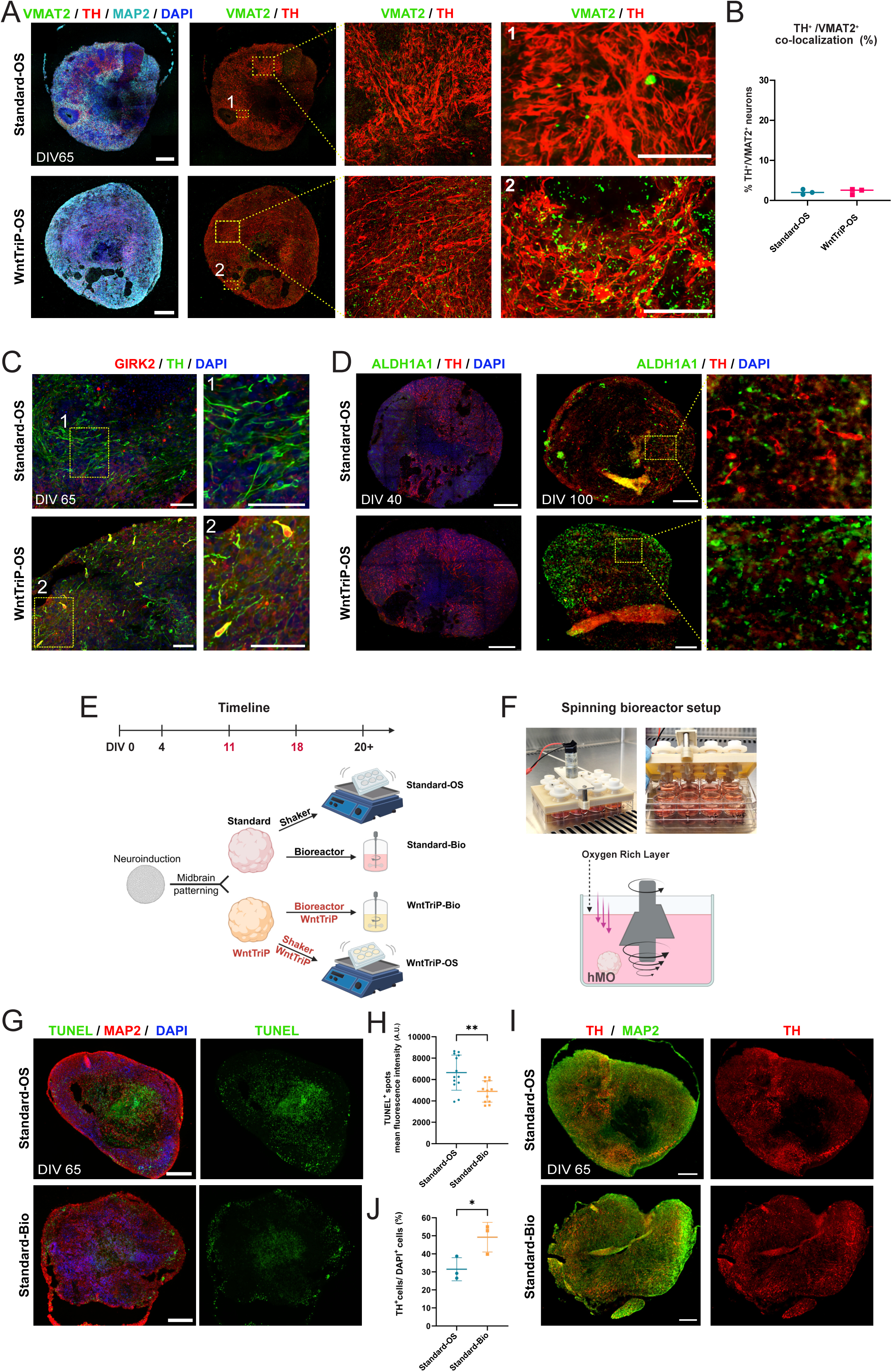
Spinning bioreactor cultivation promotes dopaminergic neuron survival and maintains tissue health in human midbrain organoids. **(A)** Representative immunofluorescence images at DIV 65 showing VMAT2 (green) and TH (red) expression in Standard-OS and WntTriP-OS organoids. High-magnification images (insets 1 and 2) are shown in adjacent panels. Scale bars: 250 µm; insets, 50 µm. **(B)** Quantification of TH^+^ spots co-localizing with VMAT2^+^ spots using the Imaris object-based spot co-localization function. Tile-stitched images from n=3 whole-hMO-sections were used. Mean ± SD. **(C)** Representative immunofluorescence images of hMO sections at DIV 65 under Standard-OS and WntTriP-OS conditions, stained for TH (green), GIRK2 (red), and nuclei (DAPI, blue). High magnification images shown in adjacent panels. Scale bars: 50 µm. **(D)** Representative images of hMO sections at DIV 40 and DIV 100 under Standard-OS and WntTriP-OS conditions, stained for ALDH1A1 (green), TH (red), and nuclei (DAPI, blue). Scale bars: 250 µm. **(E)** Schematic representation of the experimental re-design, illustrating the implementation of a bioreactor system to improve oxygen and medium exchange. **(F)** Photographs of the spinning bioreactor setup and a schematic representation illustrating the agitation technique via rotating fins. **(G)** Immunofluorescence staining for MAP2 (red) combined with a fluorometric TUNEL assay (green) to assess cell viability in organoid sections at DIV 65 under Standard-OS and Standard-Bio conditions. Cell nuclei are counterstained with DAPI. Scale bar: 250 µm. **(H)** Quantification of cell viability using the fluorometric TUNEL signal in organoid sections at DIV 65 under Standard-OS and Standard-Bio conditions. Two tile-stitched images of whole-organoid sections from two different layers were analyzed per organoid. Mean ± SD; unpaired two-tailed t-test; **p = 0.0041; n=3 organoids per condition. **(I)** Representative immunofluorescence images of TH (red) and MAP2 (green) at DIV 65 under Standard-OS and Standard-Bio conditions. Scale bars: 250 µm. **(J)** Quantification of TH^+^ neurons as a percentage of DAPI^+^ cells in organoid sections at DIV 65 under Standard-OS and Standard-Bio conditions. Mean ± SD; unpaired two-tailed t-test; *p = 0.04; n=3 organoids per condition.

To establish a more stable environment that supports long-term survival and helps maintain the newly acquired DA phenotype, we employed a spinning bioreactor^3^ to enhance oxygen and nutrient delivery during hMO differentiation **(Figure 3E, F)**. Under our culture conditions (80 rpm orbital shaking versus 100 rpm spinning bioreactor), engineering estimates derived from the Metzner-Otto relationship for stirred vessels^46^, indicate that the bioreactor provides convective mixing with lower average suspension shear (∼0.13 dyn/cm²). In contrast, orbital shaking exposes organoids to substantially higher shear (∼2.03 dyn/cm²), primarily due to friction of organoids at the well bottom. Compared to orbital shaking (Standard-OS), bioreactor-cultured organoids (Standard-Bio) exhibited significantly reduced apoptosis at DIV65, as measured by TUNEL staining (**Figure 3G,H**). Consistently, TH⁺ neuron abundance was increased under bioreactor conditions (**Figure 3I,J**). These observations were further supported at the transcriptional level by scRNA-seq. While overall cell-type composition was comparable between Standard-OS and Standard-Bio conditions (**Supplementary Figure 3B,C**), U-Cell scoring revealed reduced apoptosis signatures in bioreactor organoids, based on pro-apoptotic gene expression (e.g., *CASP3*, *BAX*, *TP53*) (**Supplementary Figure 3D,E; Supplementary Datasheet 1**). Bioreactor culture also increased expression of ion channel-related genes, including *SCN2A* and members of the *CACNA1* family, indicating enhanced electrophysiological potential, although neuronal activity and synaptic gene programs were not significantly altered (**Supplementary Figure 3F-H**). Collectively, these results demonstrate that dynamic bioreactor culture improves tissue viability and supports DA neuron survival but is not sufficient on its own to drive robust functional maturation.

### Bioreactor cultivation synergizes with tri-phasic WNT modulation to enhance DA neuron maturation and function

To evaluate whether WntTriP patterning and bioreactor cultivation synergistically enhance DA neuron maturation and stabilize the enriched SNpc-like population, hMOs were differentiated in bioreactors using either the Standard or WntTriP protocol. At DIV65, co-localization analysis of VMAT2 and TH revealed a significant increase in VMAT2 expression within DA neurons specifically in WntTriP-Bio organoids compared to all other conditions, indicating enhanced maturation of the DA compartment (**Figure 4A**). This effect was reproducible across two additional iPSC lines, with comparable astrocyte emergence across conditions, suggesting that enhanced DA maturation occurs without disrupting non-neuronal populations (**Supplementary Figure 4A,B**). Based on this clear neuronal divergence, subsequent analyses focused on direct comparisons between WntTriP-Bio and Standard-Bio organoids to isolate the contribution of tri-phasic WNT modulation within the bioreactor environment.

**Figure 4.**
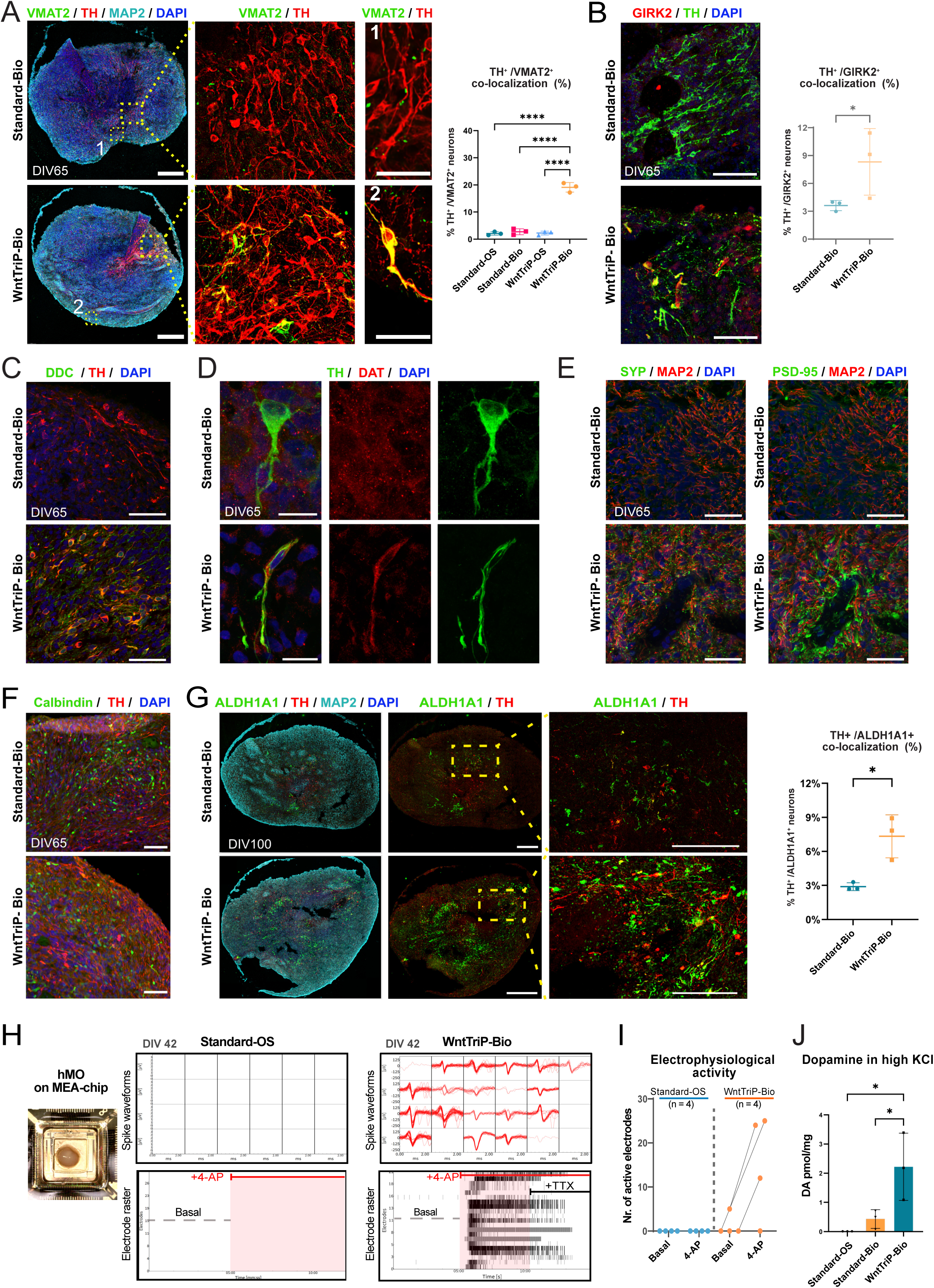
Bioreactor culture combined with tri-phasic WNT modulation enhances dopaminergic maturation, synaptic differentiation, and functional activity. **(A)** Representative immunofluorescence images of DIV65 hMOs from Standard-Bio and WntTriP-Bio conditions stained for VMAT2 (green), TH (red), MAP2 (cyan), and DAPI (blue). Higher-magnification images are shown in adjacent panels. Right: quantification of TH⁺/VMAT2⁺ co-localization across Standard-OS, Standard-Bio, WntTriP-OS, and WntTriP-Bio conditions. Mean ± SD; one-way ANOVA with Tukey’s post hoc test; ****p < 0.0001; n = 3 hMOs. **(B)** Representative immunofluorescence images of DIV65 hMOs stained for GIRK2 (red), TH (green), and DAPI (blue) under Standard-Bio and WntTriP-Bio conditions. Right: quantification of TH⁺/GIRK2⁺ co-localization. Mean ± SD; unpaired t-test; *p = 0.044; n = 3 hMOs. Scale bars: 50 µm. **(C)** Representative images of DIV65 hMOs stained for DDC (green), TH (red), and DAPI (blue). Scale bars: 50 µm. **(D)** Representative images showing TH (green), DAT (red), and DAPI (blue), indicating mature DA marker expression in WntTriP-Bio organoids. Scale bars: 15 µm. **(E)** Representative immunofluorescence images of DIV65 hMOs stained for Synaptophysin (SYP, green), PSD95 (green), MAP2 (red; pseudo color adjusted from cyan), and DAPI (blue). Scale bars: 50 µm. **(F)** Representative images of DIV65 hMOs stained for Calbindin (green), TH (red), and DAPI (blue). Scale bars: 50 µm. **(G)** Representative images of DIV100 hMOs stained for ALDH1A1 (green), TH (red), MAP2 (cyan), and DAPI (blue), with low- and high-magnification views. Right: quantification of TH⁺/ALDH1A1⁺ co-localization. Mean ± SD; unpaired t-test; *p = 0.0161; n = 3 hMOs. Scale bars: 200 µm; 50 µm. **(H)** Representative image of an hMO seeded on a 3D high-density MEA chip, with spike waveforms and electrode raster plots from DIV42 Standard-OS and WntTriP-Bio organoids under basal conditions, after 4-AP stimulation, and following TTX application. **(I)** Quantification of the number of active electrodes before and after 4-AP treatment (n = 4 organoids per condition). **(J)** Dopamine release at DIV65 measured by HPLC following high KCl stimulation in Standard-OS, Standard-Bio, and WntTriP-Bio organoids. Mean ± SD; one-way ANOVA with Tukey’s post hoc test; n = 3 hMOs per condition; *p = 0.0183 Standard-OS vs WntTriP-Bio; *p = 0.0439 Standard-Bio vs WntTriP-Bio.

Within this framework, the SNpc-associated marker GIRK2^42^ was more frequently detected in TH⁺ cells in WntTriP-Bio organoids than in Standard-Bio organoids (**Figure 4B**). Additional immunostaining showed increased expression of DA-processing markers, including DDC and DAT, in TH⁺ neurons under WntTriP-Bio conditions, consistent with a more mature DA phenotype at DIV65 (**Figure 4C,D**). Synaptic maturation was also enhanced, as indicated by increased synaptophysin and PSD95 staining. This was further supported by immunoblotting of synaptophysin normalized to βIII-tubulin (**Figure 4E**; **Supplementary Figure 5A**). To assess DA subtype specification at later stages, we examined Calbindin (CALB1), a VTA-associated marker, and ALDH1A1, associated with SNpc DA neurons ^44^. CALB1 expression at DIV65 did not differ significantly between conditions (**Figure 4F**). In contrast, at DIV100, WntTriP-Bio organoids showed a significant increase in TH⁺/ALDH1A1⁺ neurons compared to Standard-Bio, indicating improved maintenance of a mature SNpc-associated DA subtype under long-term dynamic culture conditions (**Figure 4G**). Notably, this stabilization contrasts with WntTriP-OS conditions, where TH⁺ neurons were not maintained long-term, underscoring the importance of bioreactor support. Consistently, TH⁺ neurons in WntTriP-Bio organoids displayed increased dendritic complexity at DIV100, including greater dendrite length and branching, indicative of advanced structural maturation (**Supplementary Figure 5B-D**). Given that mature DA neurons are defined not only by subtype identity but also by functional properties, we next assessed neuronal activity. HD-MEA recordings at DIV42 revealed increased numbers of active electrodes in WntTriP-Bio organoids following 4-aminopyridine (4-AP) stimulation, whereas Standard-OS organoids remained largely unresponsive at this stage (**Figure 4H,I**). Consistent with this, KCl-induced DA release at DIV65 was significantly higher in WntTriP-Bio organoids compared to both Standard-OS and Standard-Bio controls (**Figure 4J**). Together, these findings demonstrate that combining tri-phasic WNT modulation with bioreactor cultivation synergistically enhances DA neuron maturation, synaptic differentiation, electrophysiological responsiveness, and DA release, while promoting a stable SNpc-like identity.

### Single-cell transcriptomics reveals SNpc-like lineage enrichment and reduced stress signaling

To further investigate the molecular programs underlying the enhanced functionality, maturation, and VM/SNpc identity observed under WntTriP-Bio conditions, we performed scRNA-seq on DIV65 Standard-OS and WntTriP-Bio organoids, focusing on *DCX*⁺ neuronal populations. UMAP analysis resolved multiple neuronal clusters, including progenitor-like, transitional, VTA-like, mixed DA/GABAergic populations, and a terminal *SOX6*⁺/*KCNJ6*⁺ SNpc-like DA cluster (cluster 1) **(Supplementary Datasheet 1)**. This population was strongly enriched in WntTriP-Bio organoids (3.3-fold increase; 1.8 log2FC) (**Figure 5A,B**; **Supplementary Figure 6A, Supplementary Datasheet 1)** and showed robust co-expression of *EBF1*, *SV2C*, and *LMX1B*, consistent with a maturing VM and SNpc DA identity ^20, 32^ (**Supplementary Figure 6B**).

**Figure 5.**
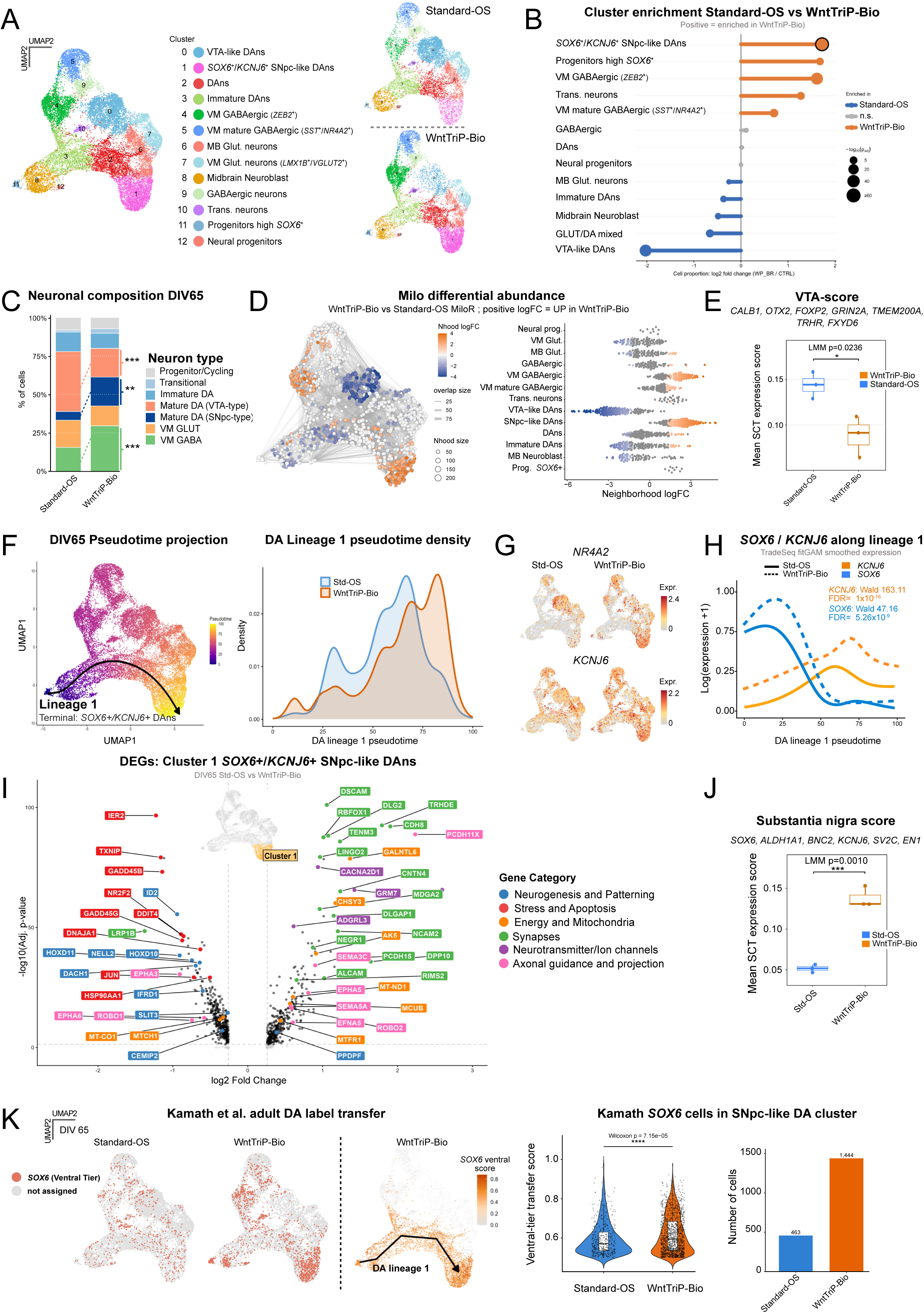
Single-cell transcriptomic analyses reveal enhanced substantia nigra-like maturation and adult ventral-tier dopaminergic features in WntTriP-Bio midbrain organoids. **(A)** UMAP visualization of the integrated DIV65 neuronal subset from Standard-OS (Std-OS) and WntTriP-Bio hMOs, colored by Seurat cluster annotation. **(B)** Differential cluster enrichment between Std-OS and WntTriP-Bio organoids at DIV65, shown as log₂ fold change in cluster proportions. Positive values indicate enrichment in WntTriP-Bio, whereas negative values indicate enrichment in Std-OS. Dot size reflects statistical significance. **(C)** Stacked bar plots summarizing neuronal composition at DIV65, highlighting ventral midbrain-associated subtypes. LMM with Benjamini-Hochberg correction; ***p = 1.05 × 10⁻LJ; **p = 1.16 × 10⁻³; ***p = 1.53 × 10⁻¹²; n = 3 hMOs per condition. **(D)** Milo neighborhood differential abundance analysis across the DIV65 neuronal manifold, showing WntTriP-Bio-enriched neighborhoods in terminal *SOX6*+/*KCNJ6*+ SNpc-like DA and VM GABAergic regions, and Standard-OS enrichment in VTA-like/immature/neuroblast-like regions.; right: beeswarm summary by cluster annotation. **(E)** VTA module score in DIV65 hMOs, showing reduced VTA identity in WntTriP-Bio organoids. LMM; *p* = 0.0236; n = 3 hMOs per condition. **(F)** Slingshot pseudotime projection onto the DIV65 UMAP with principal curves for dopaminergic lineage 1, terminating in *SOX6*⁺/*KCNJ6*⁺ SNpc-like DA neurons. **(G)** Split feature plots showing *NR4A2* and *KCNJ6* expression in Std-OS and WntTriP-Bio organoids. **(H)** TradeSeq (fitGAM) smoothed expression dynamics of *SOX6* and *KCNJ6* along lineage 1 pseudotime. **(I)** Differential gene expression in cluster 1 (*SOX6*⁺/*KCNJ6*⁺ SNpc-like DA neurons) between Std-OS and WntTriP-Bio organoids at DIV65; colored annotations indicate functional gene categories. **(J)** Substantia nigra module score in cluster 1 (genes: *SOX6, KCNJ6, ALDH1A1, SV2C, EN1, BNC2*), showing increased scores in WntTriP-Bio. LMM; ***p = 0.0010; n = 3 hMOs per condition. **(K)** Label transfer using the Kamath et al. dataset projected onto the DIV65 neuronal subset, classifying cells into *SOX6*⁺ ventral-tier identities. Left: projection of Kamath *SOX6* scores along dopaminergic lineage 1. Right: violin plot of ventral-tier scores in cluster 1. Wilcoxon test; ****p = 7.15 × 10⁻LJ.

At the level of neuronal subtype composition, WntTriP-Bio organoids displayed a significant reduction in VTA-like DA neurons, accompanied by an increase in mature VM populations, including VM GABAergic neurons and a 3.6-fold expansion of *SOX6*⁺/*KCNJ6*⁺ SNpc-like DA neurons by DIV65 (**Figure 5C; Supplementary Datasheet 1**). To capture these shifts beyond discrete clustering, we applied Milo neighborhood differential abundance analysis across the neuronal manifold. This revealed that WntTriP-Bio hMOs were strongly enriched for *SOX6*^+^/*KCNJ6*^+^ SNpc-like DA neighborhoods and VM GABAergic neighborhoods, while VTA-like DA neurons, immature DA neurons, and midbrain neuroblast neighborhoods were depleted relative to Standard-OS hMOs, indicating a more mature and nigral-like VM cellular composition. **(Figure 5D; Supplementary Datasheet 1**). Consistent with this strong shift from VTA-like to SNpc-like DA neurons, marker analysis across neuronal populations showed reduced expression of VTA-associated genes and increased expression of SNpc-associated markers in WntTriP-Bio organoids relative to Standard-OS (**Supplementary Figure 6C**). Accordingly, a composite VTA gene score was significantly decreased across the neuronal compartment (**Figure 5E, Supplementary Datasheet 1**). Together, these results indicate that WntTriP-Bio reshapes the neuronal landscape toward a more mature, nigral-like configuration, while Standard-OS organoids retain a comparatively immature and VTA-biased profile.

To further examine lineage progression, we reconstructed DA differentiation trajectories at DIV65 (**Figure 5F**). Pseudotime analysis identified a trajectory terminating in the *SOX6*⁺/*KCNJ6*⁺ SNpc-like cluster, with WntTriP-Bio organoids showing increased enrichment at these terminal states, indicative of enhanced maturation toward an SNpc-like fate. Condition-specific feature plots for *NR4A2* and *KCNJ6*, together with tradeSeq (fitGAM) analysis of *SOX6* and *KCNJ6* along lineage 1, indicated that WntTriP-Bio organoids sustain higher expression of SNpc-associated genes while maintaining early *SOX6*-driven specification programs (**Figure 5G,H**). We next assessed whether these terminal SNpc-like neurons exhibit distinct functional transcriptional programs. Differential expression analysis identified 377 upregulated and 421 downregulated genes in WntTriP-Bio organoids (**Figure 5I, Supplementary Datasheet 1**). Upregulated genes were enriched for pathways related to synaptic organization, neurotransmission, ion channel activity, axonal growth, and mitochondrial metabolism, whereas genes associated with stress responses, apoptosis, and immature neurogenic programs were reduced compared to Standard-OS (**Figure 5I**; **Supplementary Figure 6C,D**). In line with these findings, an SNpc gene module score (*SOX6*, *ALDH1A1*, *BNC2*, *KCNJ6*, *SV2C*, *EN1*) was significantly increased in cluster 1 under WntTriP-Bio conditions, with high consistency across biological replicates (**Figure 5J**). Notably, pySCENIC analysis identified *EBF1* regulon activity as the most strongly upregulated in WntTriP-Bio organoids, suggesting a potential central role for *EBF1* in driving SNpc-like maturation and identity (**Supplementary Figure 6E, Supplementary Datasheet 1**). In contrast, regulons enriched in Standard-OS organoids were dominated by immature neurogenic (e.g., *SOX2*) and stress- or apoptosis-related programs (e.g., *CREB3*, *DDIT3*) ^47, 48^. Consistent with this, biological process enrichment analysis in the SNpc-like DA cluster revealed significant enrichment of pathways related to synapse organization, trans-synaptic signaling, neurotransmission, neuron migration, and axon development under WntTriP-Bio conditions (**Supplementary Figure 6F**), further supporting a link between EBF1 activity and maturation of SNpc-like DA neurons. Finally, label transfer using an adult human DA reference by Kamath et al.^22^ showed that WntTriP-Bio organoids were shifted toward *SOX6*⁺ ventral-tier identities, with increased accumulation of these cells at the terminal stage of lineage 1 pseudotime (**Figure 5K**), corresponding to DA populations known to be particularly vulnerable in PD. These findings demonstrate that WntTriP bioreactor culture not only promotes progression toward a late-stage, disease-relevant SNpc-like DA identity, but also enhances neuronal maturation through strengthened synaptic and metabolic programs while reducing stress- and apoptosis-associated transcriptional signatures.

### α-synuclein preformed fibrils induce DA neuron loss and proteinase K–resistant aggregates in WntTriP bioreactor derived human midbrain organoids

To validate the WntTriP-Bio iPSC-derived VM hMOs as a tool to model PD pathology, we exposed the organoids to α-synuclein preformed fibrils (αSyn-PFFs) and evaluated the resulting effects over time **(Figure 6A)**. To confirm αSyn-PFF uptake, we first utilized Alexa-594 labelled αSyn-PFFs (α-Syn-PFF 594). Immunofluorescence analysis revealed an efficient uptake of αSyn-PFF 594 within 5 days of treatment, with a high density observed in the outer layer of organoids **(Figure 6B, C)**. Next, we employed unlabelled αSyn-PFFs to eliminate any potential off-target effects of the labeling dye^49^. Immunofluorescence analysis of αSyn-PFF-treated organoids revealed comparable TH^+^ neuronal density at DIV 85 (5 days post-treatment) in both Tris saline-treated (control) and αSyn-PFF treated organoids **(Figure 6D)**. However, a significant reduction of TH^+^ neurons was observed in αSyn-PFF-treated organoids 30 days post-treatment, highlighting the vulnerability of DA neurons to αSyn-PFFs **(Figure 6E, F)**. Next, to assess pathological aggregates of αSyn, we performed immunostaining for αSyn **(Figure 6G)** with and without proteinase K treatment^50^. Following proteinase K digestion of DIV 110 organoid sections **(Figure 6H)**, only the αSyn-PFF treated organoids displayed proteinase K-resistant α-Syn deposits. These deposits exhibited a shape indicative of localization in neuronal cells **(Figure 6H)**. In contrast, controls showed minimal to no proteinase K-resistant αSyn signal, indicating the absence of pathological α-Syn aggregates. Taken together, these findings demonstrate that the WntTriP-bioreactor midbrain organoid model, following αSyn-PFF exposure, effectively recapitulates key pathological hallmarks of PD, including the loss of DA neurons and the formation of insoluble αSyn aggregates.

**Figure 6.**
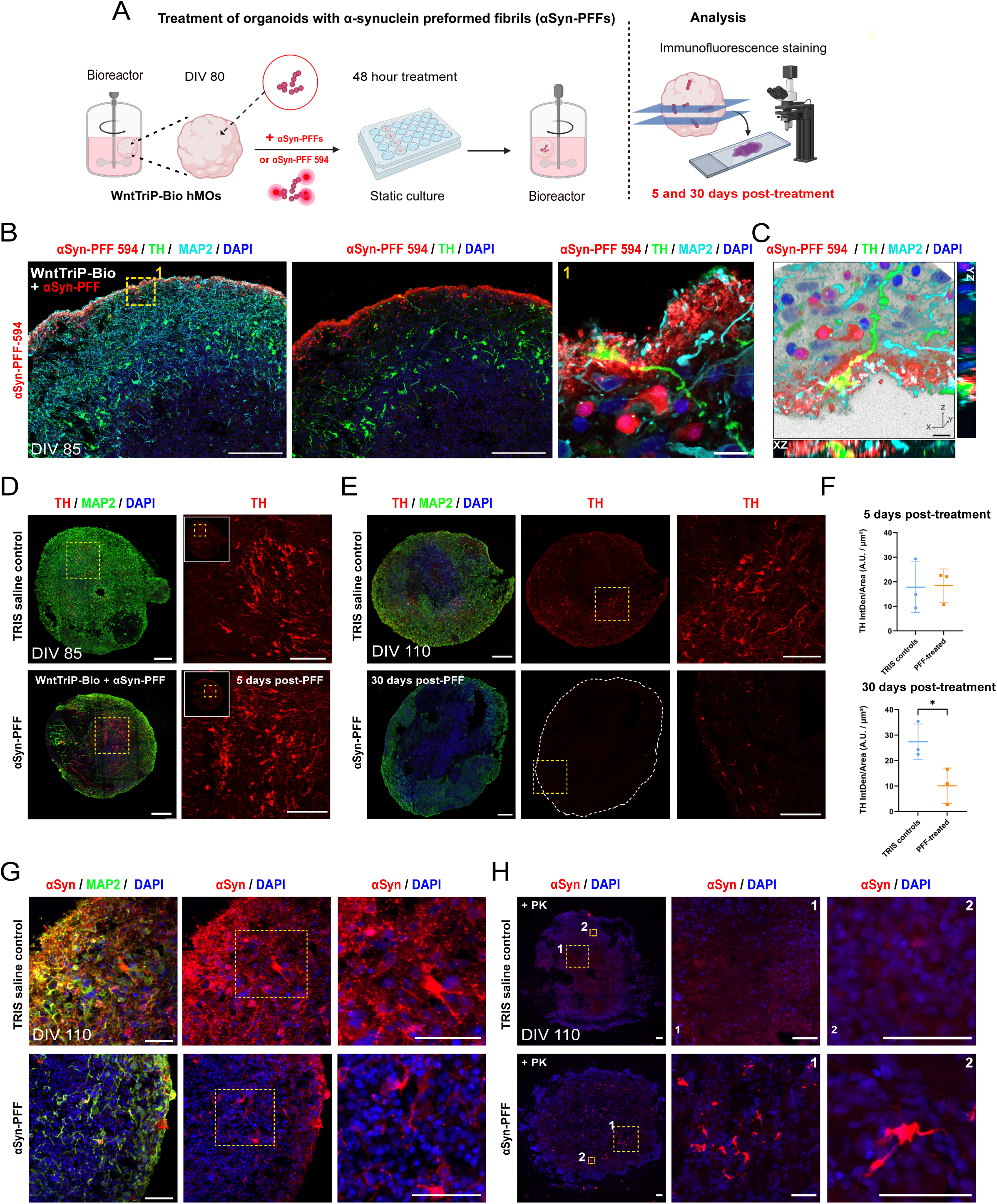
α-synuclein preformed fibrils induce dopaminergic (DA) neuron loss and proteinase K-resistant deposits in WntTriP bioreactor midbrain organoids. **(A)** Schematic diagram illustrating the experimental design for α-synuclein preformed fibrils (αSyn-PFF) treatment of WntTriP-Bio organoids, including treatment timeline and analysis endpoints. **(B)** Representative images of WntTriP-Bio hMO at DIV 85 (5 days post-treatment with labelled αSyn-PFF 594) showing αSyn-PFF 594 in red, TH (green) and MAP2 (cyan) to display internalization. Boxed region 1 is shown at higher magnification in the adjacent panel. The subsequent panels D-G depict organoids treated with unlabelled αSyn-PFFs. **(C)** 3D-reconstruction of confocal z-stack images, showing labelled αSyn-PFF 594 (red) neuronal uptake together with TH, (green), MAP2 (cyan). Orthogonal slices of the 3D-volume are shown in the adjacent panels below (XZ orientation) and to the right (YZ orientation). Scale bar: 15 µm. **(D)** Immunofluorescence images of hMO at DIV 85 (5 days after αSyn-PFF treatment) showing TH^+^ neurons in TRIS-Saline-treated controls (top) and αSyn-PFF treated organoids (bottom). Sections were labelled against TH (red) and MAP2 (green). Cell nuclei were counterstained with DAPI. Scale bars: 200 μm in tile-stitched organoid sections, 100 μm in zoom-ins. **(E)** HMOs at DIV 110 (30 days after αSyn-PFF treatment) show a loss of TH-positive DA neurons in αSyn-PFF treated organoids (bottom) compared to TRIS-Saline controls (top). Sections were labelled against TH (red) and MAP2 (green). Cell nuclei were counterstained with DAPI. Scale bars: 200 μm in tile-stitched organoid sections, 100 μm in zoom-ins. **(F)** Quantification of TH^+^ DA neurons in TRIS-Saline controls and αSyn-PFF treated WntTriP-Bio organoids, 5 days after treatment (DIV85) and 30 days after treatment (DIV110). Mean ± SD; unpaired two-tailed t-test. *p = 0.0386; Tile-stitched images of whole-organoid-sections were used; n=3 organoids per condition. **(G)** Immunostaining for MAP2 (green), αSyn (red) and DAPI (blue) in organoid sections +/- PFF, without proteinase K (PK) treatment (control for panel H). Boxed regions are shown at higher magnification in the adjacent panels. Scale bars: 50 μm. **(H)** Immunostaining for αSyn (red) and DAPI (blue) in organoid sections treated with PK reveals protease-resistant αSyn aggregates in samples exposed to αSyn-PFFs. Boxed regions 1 and 2 are shown at higher magnification in the adjacent panels. Scale bars: 200LJμm (tile-stitched sections); 50LJμm (zoomed-in views).

## Discussion

In this study, we established an integrated strategy to generate hMOs enriched in functionally mature DA neurons exhibiting molecular, morphological, and physiological features consistent with mDA identity. By combining developmentally informed signaling cues with scalable, physiologically supportive culture conditions, we developed a robust and reproducible platform for modeling mDA neuron vulnerability and PD-associated neurodegeneration. Beyond recapitulating key aspects of regional DA neuron identity, this system provides a versatile and translationally relevant platform for investigating disease mechanisms and screening candidate therapeutics.

Modeling PD using iPSC-derived systems remains challenging due to the limited generation of regionally specified, vulnerable DA neuron subtypes and the difficulty in reproducing hallmark pathological features such as protein aggregation and progressive neurodegeneration. In the human brain, PD pathology predominantly affects DA neurons in the ventral tier of the SNpc ^11, 51^, characterized by markers such as SOX6, ALDH1A1, and GIRK2 ^21^. While most of our current understanding derives from rodent models, significant interspecies differences distinguish DA neurons in their quantity, organization, connectivity, molecular identity and developmental origin. Comparative single-cell transcriptomic analyses have highlighted species-specific differences in vulnerable DA subtypes, with key markers such as SOX6, ALDH1A1, and GIRK2 being absent, underrepresented, or differently expressed in rodent SNpc^21, 22^. In addition, αSyn biology differs substantially between species in terms of aggregation propensity, post-translational modification, and propagation behavior ^52–54^. These discrepancies limit the translational relevance of rodent models and underscore the need for human-specific systems such as hMOs.

In this context, hMOs represent a promising platform to recapitulate human midbrain tissue *in vitro*, and substantial efforts have been made to improve DA neuron specification and yield ^55–57^. Nevertheless, current models often fail to generate sufficiently mature and functionally competent neurons, and particularly struggle to maintain DA populations in long-term culture, which is an essential requirement for modeling neurodegenerative processes.

Here, we addressed these limitations by combining tri-phasic WNT modulation with bioreactor-based culture. At early stages, WntTriP treatment (DIV25) directed organoids toward a midbrain-restricted, ventral/SNpc-biased transcriptional state, characterized by *SOX6*⁺/*EBF1*⁺ populations, increased VM/SNpc gene signatures, and accelerated DA lineage progression. At later stages, integration with bioreactor culture promoted the emergence of a terminal *SOX6*⁺/*KCNJ6*⁺ SNpc-like DA population with strong *EBF1* co-expression and features consistent with adult ventral-tier identity. These transcriptional changes were accompanied by increased expression of maturation and functional markers, including VMAT2, GIRK2, DAT, and synaptophysin, as well as enhanced electrophysiological activity and DA release, indicating coordinated subtype specification and functional maturation.

The critical role of WNT signaling in DA lineage specification is well established, with precise temporal modulation required to support progenitor expansion and differentiation^15, 16, 58^. Consistent with prior studies in 2D systems^59^, our results demonstrate that temporally controlled WNT inhibition following progenitor priming enhances neuronal output and refines VM identity. WntTriP organoids maintained strong regional patterning while promoting progression toward a *SOX6*⁺/*KCNJ6*⁺/*EBF1*⁺ lineage, accompanied by a coordinated upregulation of numerous synaptogenic programs. Pseudotime, tradeSeq, and regulatory network analyses further revealed accelerated and coordinated activation of key SNpc-associated genes, including *EN1*, *SV2C*, *SOX6*, *KCNJ6*, *EBF1*, *SLIT2*, and *ERBB4*. Notably, *EBF1* expression peaked in the most mature DA cluster, consistent with its role in nigral DA neuron maturation^20^, suggesting faithful specification of SNpc-like identity. Importantly, this enhanced molecular maturation translated into earlier and increased functional activity. WntTriP-Bio organoids displayed neuronal responsiveness as early as DIV42 and exhibited significantly higher DA release compared to controls. These effects were accompanied by reduced expression of stress- and apoptosis-associated pathways, indicating that improved maturation occurs in a more supportive cellular environment. This relationship between reduced cellular stress and enhanced maturation may be particularly relevant, or even necessary, for enabling the emergence of SNpc DA neurons, whose high bioenergetic demands contribute to their selective vulnerability in PD ^22, 26^. The contribution of the bioreactor environment was therefore critical in this context.

*In vivo*, the primate SNpc exhibits relatively dense vascularization compared to other midbrain regions, a feature that may be associated with its elevated bioenergetic demands and heightened vulnerability to degeneration^60^. In contrast to the extensive vascularization and oxygenation *in vivo*, conventional *in vitro* systems often suffer from limited oxygen and nutrient diffusion, particularly in larger organoids ^61, 62^. To compensate for these limitations, bioreactor culture improves convective mixing, enhances oxygenation, and reduces shear stress compared to orbital shaking, thereby supporting tissue viability and metabolic homeostasis. Under our conditions, reduced mechanical stress combined with improved oxygen and nutrient delivery likely contributed to the enhanced survival, maturation, and stability of DA neurons.

Notably, projection onto adult human midbrain reference datasets ^22^ revealed that WntTriP-Bio organoids are enriched in *SOX6*⁺ DA populations corresponding to ventral-tier neurons most vulnerable in PD. In addition, this approach promoted the expansion of VM GABAergic populations, which are known to reside in the VM and contribute to local circuit regulation of DA neuron activity ^30, 63^. These findings suggest that WntTriP-Bio not only enhances SNpc-like DA identity but also supports the formation of a more physiologically relevant cellular niche.

WntTriP-Bio organoids generate a larger population of mature, functionally active DA neurons compared to standard conditions. In humans, SNpc DA neurons are particularly susceptible to metabolic stress due to their extensive axonal arborization, dense synaptic connectivity, and intrinsic pacemaking activity, all of which impose high bioenergetic demands and contribute to their vulnerability in aging and disease ^26, 64–66^. In this context, neuronal activity and synaptic dysfunctions, particularly at presynaptic terminals where αSyn is predominantly active, have been closely linked to PD pathogenesis ^67,68^. Consistent with this, exposure to αSyn-PFFs resulted in a pronounced loss of DA neurons in WntTriP-Bio organoids, likely reflecting their advanced maturation and functional state, including the focused upregulation of synaptogenic programs. Indeed, synaptic activity has been closely linked to pathological αSyn phosphorylation and aggregation ^69, 70^. Accordingly, WntTriP-Bio organoids showed accumulation of proteinase K-resistant αSyn following PFF exposure, supporting their utility for modeling synucleinopathy-related neurodegeneration.

In conclusion, we demonstrate that combining tri-phasic WNT modulation with dynamic bioreactor culture synergistically enhances the specification, maturation, and functional integration of SNpc-like DA neurons in hMOs. This approach overcomes key limitations of existing models by enabling both early VM patterning and long-term maintenance of mature, functionally active DA neurons. Beyond PD, this platform is relevant for modeling neuropsychiatric disorders, as dysregulation of DA signaling, mitochondrial function, and synaptic activity represents a shared pathogenic axis across multiple conditions ^71–73^. By recapitulating human DA neuron identity, activity, and selective vulnerability, WntTriP bioreactor-derived hMOs provide a robust and translationally relevant system for mechanistic studies and the development of predictive human disease models.

## Methods

### Ethics approval and consent to participate for experiments involving human samples

All cells used in the study were derived from patients who signed an informed consent form, approved by the ethics committee of the Medical Faculty at the University Hospital Tübingen (Ethikkommission der Medizinischen Fakultät am Universitätsklinikum Tübingen). No identifiable images of human research participants are included. All experiments and protocols were approved by the Comité d’Évaluation Éthique de l’Inserm / Inserm Institutional Review Board (CEEI/IRB; IRB00003888; reference CD/EB 24-002). All experiments were conducted in accordance with the relevant institutional and ethical guidelines and regulations.

### Human induced pluripotent stem cell culture

The following human induced pluripotent stem cell (iPSC) lines were used in the study: Kolf2.1J (RRID:CVCL_B5P3, Jackson laboratory, Bar Harbor, USA) and two previously generated and characterized healthy control lines, labelled as C2 and C3^74, 75^. Human iPSCs were grown on Matrigel-coated (Corning, Cat # 354230, New York, USA) dishes in mTeSR1 medium (StemCell Technologies, Cat # 85850, Vancouver, Canada) supplemented with 1% penicillin-streptomycin solution (ThermoFisher, Cat # 15140122, Waltham, MA, USA). The iPSC medium was changed every other day. Cells were passaged using ReLeSR dissociation agent (StemCell Technologies, Cat # 100-0483) every 5-7 days at a 1:12 ratio to maintain optimal growth and pluripotency. IPSCs were routinely tested negative for mycoplasma using the Venor®GeM Classic Mycoplasma detection kit (Minerva-Biolabs, Cat # 11-1025G, Berlin, Germany) and kept below passage 35.

### Generation of human midbrain organoids

Midbrain organoids of the “Standard” condition were generated according to a protocol previously described^7^. IPSCs were dissociated into single cells using Accutase (StemCell Technologies, Cat # 07920, Vancouver, Canada) and seeded into low-attachment U-bottom 96-well plates (Corning, Cat # 7007, New York, USA) at a density of 9,000 cells per well in a 1:1 mix of DMEM F12 (Gibco, Cat #12634010) and Neurobasal medium (Gibco, Cat # 21103049) supplemented with 0.5% N-2 supplement (Gibco, Cat # 11520536), 1% B27 supplement without vitamin A (Gibco, Cat # 12587010), 1% NEAA (Gibco, Cat # 11140050), 0.01% β-mercaptoethanol, 1 µg/mL heparin (Merck, Cat # H3149-10KU), 10 µM SB431542 (Tocris, Cat # 1614), 200 ng/mL LDN (Tocris, Cat # 6053), 0.7 µM CHIR99021 (Sigma-Aldrich, Cat # SML1046), and 50 µM Rock inhibitor Y-27632 (Tocris, Cat # 1254). On day 4, the medium was changed to DMEM F12/Neurobasal (1:1) supplemented with 0.5% N2 supplement, 1% B27 supplement, 1% NEAA, 0.1% β-mercaptoethanol, 1 µg/mL heparin, 10 µM SB431542, 200LJng/mL LDN, 100 ng/mL SHH-C25II (PeproTech, Cat # 100-45), 1 µM purmorphamine (Sigma, Cat # 540220), 7.5 µM CHIR99021, and 100 ng/mL FGF8 (PeproTech, Cat # 100-25). On day 7, embryoid bodies (EBs) were embedded in Matrigel. Matrigel was diluted in a 3:2 ratio with medium and used as an embedding mixture. EBs were washed in fresh medium and embedded in Matrigel in a six-well ultralow-attachment plate using the Matrigel-medium mixture. The Matrigel-EB mixture was incubated for 30 min at 37LJ°C, and on day 7, medium containing the following compounds was added: Neurobasal medium supplemented with 0.5% N2 supplement, 1% B27 without Vitamin A supplement, 1% GlutaMax (Gibco, Cat # 35050061), 1% NEAA, 0.1% β-mercaptoethanol, 2.5 µg/mL insulin (Gibco, Cat # 12585014), 200 ng/mL laminin (Thermo Fisher, Cat # 23017015), 7.5 µM CHIR99021, 100 ng/ml SHH-C25II, 1 µM purmorphamine, and 100 ng/mL FGF8. On day 8, the medium was changed to Neurobasal medium supplemented with 0.5% N2, 1% B27, 1% GlutaMax, 1% NEAA, 7.5 µM CHIR99021, 0.1% β-mercaptoethanol, 10 ng/mL BDNF (PeproTech, Cat # 450-02), 10 ng/mL GDNF (Peprotech, Cat # 450-10), 100 µM ascorbic acid (Sigma, Cat # A8960), and 125 µM dbcAMP (MedChem, Cat # HY-B0764). On day 10, the medium was refreshed, and the concentration of CHIR99021 was reduced to 3 µM. On day 13, the concentration of CHIR99021 was further reduced to 0.7 µM until day 15. On day 20, the Matrigel was dissociated from the organoids and the organoid-containing well-plates were transferred to an orbital shaker inside the incubator (Heracell^TM^ 240i CO_2_ Incubator, ThermoFisher, Cat # 16426639, Waltham, MA, USA). For the bioreactor-cultivation method, organoids were transferred to a multi-well spinning bioreactor at DIV 20, after dissociation from the Matrigel. The spinning bioreactor was purchased from 3Dnamics (Customized SpinΩ Bioreactor – 12-well, Cat # 3DNA01, Baltimore, MD, USA) and its design is based on the initial publication of Qian et al., 2018^3^. The 3D-printed ULTEM plastic, motor-driven spinning module serves as the lid for conventional 12-well low-attachment plates. It features 12 angled rotator shafts that independently agitate the medium in each well.

For the WNT signaling inhibition condition, we adapted and modified a recent strategy by inhibiting WNT signaling after midbrain floor plate induction^15^. Differentiation medium was supplemented with the small molecule WNT/β-catenin signaling inhibitor IWP-2 (STEMCELL, Cat #72122) at a concentration of 2.2 μM from day 11 to day 18, refreshed every other day. In this condition, the WNT/β-catenin activator CHIR99021 was completely removed from the medium starting on day 11. All other media components remained identical to the standard condition. From day 18 onwards, the medium for all conditions was replaced with the final maturation medium, consisting of Neurobasal medium supplemented with 0.5% N2, 1% B27 without Vitamin A, 1% GlutaMax, 1% NEAA, 0.1% β-mercaptoethanol, 10 ng/mL BDNF, 10 ng/mL GDNF, 100 µM ascorbic acid, and 125 µM dbcAMP. On day 20, in all conditions, Matrigel was mechanically dissociated from the organoids, and the well plates containing organoids were transferred to either an orbital shaker or a spinning bioreactor inside the incubator.

### Immunohistochemistry and imaging

Organoids were fixed in 4% paraformaldehyde (PFA, ROTI Histofix; Roth, Cat # P087.1) for 1 hour at room temperature, washed three times in PBS (5 minutes each), cryoprotected in 30% sucrose at 4°C overnight, embedded in optimal cutting temperature compound (OCT, Tissue-Tek, Cat # 0094-4583-01), and sectioned on Superfrost PLUS tissue adhesion glass slides (Epredia, Cat # J1800AMNZ, Kalamazoo, MI, USA) at 20 μm using a Leica CM3050S cryostat (Leica). For immunohistochemistry, sections were air-dried for 10 minutes followed by sodium citrate buffer (pH 6.0) guided antigen retrieval for 20 minutes at 85°C. After cooling to room temperature, sections were blocked for one hour with 10% normal donkey or goat serum in PBS containing 0.2% Triton X-100 (Sigma, Cat # X100-5ML) and incubated overnight at 4°C with primary antibodies. Primary and secondary antibody lists are provided in **Supplementary Table 2**. After primary antibody incubation, sections were washed three times with PBS and then incubated for 2 hours at room temperature with species-specific secondary antibodies. This was followed by three 5-minute washes in PBS. Nuclei were counterstained with 4′,6-diamidino-2-phenylindole (DAPI) at a concentration of 1 µM for 5 minutes and mounted with DAKO (Agilent DAKO, Cat # S302380-2, Glostrup Municipality, Denmark) mounting medium for image acquisition. For experiments implementing proteinase K, sections were treated with proteinase K (5 μg/mL, 10 minutes at room temperature) to digest soluble proteins while preserving insoluble aggregates. Control sections were handled the same way but without adding proteinase K to the incubation solution. Both groups were then immunostained with primary antibodies.

For confocal imaging, whole-section organoid confocal tile scan images were acquired using a Leica TCS SP8 confocal microscope (Leica, Germany) equipped with a 40xLJ/1.4 numerical aperture oil-immersion objective. Tile-scanned fluorescence microscope images were stitched using the automated stitching function of the Leica LasX software package (10% tile overlap; Smooth stitch option). For high magnification imaging the 63x/1.4 NA objective was used. Images were analyzed using ImageJ or Imaris software.

### Fluorometric TUNEL assay and imaging

Organoids were fixed in 4% PFA for 1 hour at room temperature, cryoprotected in 30% sucrose overnight, embedded in OCT, and sectioned at 20 μm using a cryostat. Sections were blocked for one hour with 10% normal goat serum in PBS containing 0.2% Triton X-100 and incubated overnight at 4°C with primary antibodies. Following incubation with primary antibodies, sections were washed three times in PBS and incubated with species-specific secondary antibodies conjugated to Alexa Fluor 568 and Alexa Fluor 647 dyes (1:1000, Invitrogen, Cat # A-11011; Cat # A48260, Waltham, MA, USA) for 1 hour at room temperature, followed by three washes of 5 minutes each in PBS. Apoptotic nuclei were detected using the DeadEnd™ Fluorometric TUNEL System, following the manufacturer’s instructions (Promega, Cat # G23250, Madison, WI, USA). Sections were subjected to 100 µL equilibration buffer (compounds supplied by the DeadEnd^TM^ TUNEL kit) for 8 minutes at room temperature followed by labeling with 50 µL of the TdT reaction mix for 60 minutes at 37°C in a humidified chamber. To stop the reaction, sections were immersed in 2X Saline Sodium Citrate buffer (2XSSC) for 15 minutes at room temperature followed by three washes in PBS, 5 minutes each. Nuclei were counterstained with DAPI and mounted with DAKO mounting medium for image acquisition. Whole-section organoid confocal tile scan images were acquired using a Leica TCS SP8 confocal microscope (Leica, Germany) equipped with a 40xLJ/1.4 numerical aperture oil-immersion objective. Images were pre-processed using ImageJ software and then analyzed using Imaris software (version 9.2) through the “Spots Intensity” function. TUNEL^+^ spots were detected across 12 organoid sections from different layers in n=4 organoids per condition. Mean TUNEL^+^ spot intensities were calculated per organoid and compared. Overall detected TUNEL^+^ spots were 25913 in the bioreactor condition and 56065 spots in the standard condition.

### Quantitative analysis of DA neurons in midbrain organoids

To quantify DA neurons, organoid sections were stained for TH and the number of TH^+^ neurons was counted across entire sections from at least three independent organoids per condition. Image processing was performed using ImageJ software. Data were normalized to the total number of cells (DAPI^+^ nuclei) and expressed as a percentage.

### Measurement of DA neuronal branching

Neuronal branching was quantified using the Filament Tracer module in Imaris (version 9.2, Bitplane, Belfast, UK). Confocal z-stacks of TH-stained organoid sections were imported into the software for 3D reconstruction of DA neuron structures. Parameters were set to detect soma, dendrites, and axons. Quantified metrics included the total number of DA somas, total filament and dendrite length, and the number of branching segments per filament to assess neuronal arborization. Representative images were processed in ImageJ.

### ScRNA-seq - transcriptome library preparation and sequencing

IPSC-derived midbrain organoids were dissociated into single-cell suspensions using a Worthington Papain Dissociation System kit (Worthington Biochemical, Cat # LK003153), according to manufacturer’s instructions. Individual organoids were minced and incubated with 2.5LJmL papain/DNase solution on an orbital shaker for 30 min at 37 °C. Organoid suspensions were minced using scalpels and 1 mL low-attachment pipette tips and then incubated on an orbital shaker for 20 min at 37 °C twice. After enzyme inactivation, the cell suspensions were filtered through a 40-μm cell strainer. The final cell suspension was centrifuged at 300×g for 7 min, and the cell pellet was resuspended in PBS-0.04% BSA at a final concentration of 1000 cells/μL, ensuring that more than 95% of the cells were viable. scRNA-Seq libraries were generated with the 10X Chromium Next GEM Single Cell 3’ Reagent Kit v3.1 (Cat. number 1000128, 10x Genomics, Pleasanton, CA, USA) according to the manufacturer’s instructions. Libraries were pooled and subjected to paired-end sequencing on the Illumina NovaSeq X platform (SP Flow Cell, Illumina) with a sequencing depth of 300 million reads per library at GENEWIZ GmbH (Leipzig, Germany). The data have been deposited into GEO under accession number GSE301365.

### ScRNA-seq data analysis

The sequencing data were demultiplexed and filtered via the 10X Genomics Cell Ranger pipeline to generate filtered gene-barcode matrices, which were used as input for downstream analysis with Seurat (v5.3.0) and R (v4.4.1)^76, 77^. DoubletFinder was used to detect doublets and SCTransform was used for normalization while regressing cell-cycle effects. Datasets were integrated using Harmony, followed by principal component analysis (PCA), nearest-neighbor graph construction, clustering and UMAP for visualization. Marker genes were identified using FindAllMarkers, and differential expression between conditions within each cluster was computed using FindMarkers with the MAST test and Bonferroni correction. For focused DIV65 analyses, a pre-integrated *DCX*^+^ neuronal subset, subsampled to equal cell numbers between conditions, was analyzed separately from the full object. Sample-aware comparisons of module scores, identity scores, label-transfer projections, and pseudotime-derived metrics were performed using linear mixed models or generalized linear mixed models, with sample included as a random effect.

### Trajectory analysis using Slingshot and tradeSeq

Trajectory inference was performed with Slingshot (v2.12.0)^78^. For DIV25 analyses, FPP populations were defined as the developmental root, and DA-associated terminal clusters were used to infer lineage trajectories. For DIV65 neuronal analyses, with progenitors (cluster 11) set as the root, lineage 1 was defined across clusters 11-8-3-2-1. The inferred trajectory was projected onto the neuronal UMAP to compare lineage occupancy between conditions.

Condition-dependent gene dynamics along pseudotime were modelled using tradeSeq (fitGAM) (v1.18.0)^79^. Raw count data from cells assigned to the SOX6⁺ DA Slingshot lineage were fitted using negative binomial generalized additive models (GAMs) via *fitGAM* (6 knots), incorporating parallel condition-specific trajectories. Differential expressions across pseudotime between conditions was assessed using *conditionTest*, which applies a Wald test to compare GAM coefficients at each knot. P-values were adjusted for multiple testing using the Benjamini–Hochberg procedure (FDR < 0.05). Smoothed expression curves were generated by interpolating predictions onto a common pseudotime grid and averaging across lineages within each condition.

### Milo differential abundance analysis

Condition-dependent changes in neighborhood abundance across the single-cell manifold were assessed using miloR (v2.0.0)^80^ on the Harmony-integrated Seurat objects. Seurat objects were converted to SingleCellExperiment (v1.26.0), neighborhood graphs were constructed on the first 30 Harmony dimensions using buildGraph and makeNhoods (k = 30, prop = 0.08, refined = TRUE), and sample-level neighborhood counts were tested with testNhoods using the design ∼ condition. Differential abundance was summarized as neighborhood logFC, and significance was defined at SpatialFDR < 0.10 and significant neighborhoods were then visualized by neighborhood logFC on the Milo graph and beeswarm summaries. Significant Milo neighborhoods were projected back onto the UMAP and summarized by final cluster annotation.

### Gene and module scores

Gene set and identity scores were calculated on normalized SCT expression data as the per-cell mean expression of the genes in each predefined signature, followed by condition- or sample-level summarization for visualization and statistical testing. The identity and module scores including gene-sets used in the manuscript are listed in the **Supplementary Datasheet 1.**

### pySCENIC analysis

Regulon activity in the DIV65 neuronal subset was inferred using pySCENIC (v0.12.1)^81^. Raw count matrices exported from Seurat were converted from matrix market format to loom format with loompy (v3.0.8; Python 3.6). Gene regulatory networks were first inferred with pySCENIC GRN using the human transcription factor list, followed by motif enrichment and regulon pruning with pySCENIC CTX using the cisTarget ranking database and motif annotation table. Regulon activity was then scored in individual cells with AUCell (v1.32.0), and the resulting AUC matrix was imported back into Seurat as a dedicated assay for downstream visualization and differential regulon activity analysis.

### CellChat analysis

Cell-cell communication at DIV25 was inferred using CellChat v2.2.0^82^ on the group-balanced DIV25 single-cell dataset (21,040 Standard-OS and 21,040 WntTriP-OS cells). Condition-specific CellChat objects were generated across nine annotated cell populations, communication probabilities were computed using the triMean method, interactions supported by fewer than ten cells in either sender or receiver populations were removed, and pathway-level networks were merged for comparative analysis. Differential ligand-receptor interactions involving SOX6^+^ DA neurons were then examined within VM-relevant morphogen, axon-guidance, adhesion, and synaptic pathway families to define the signaling niche associated with the emerging SNpc-like DA state.

### Dopamine measurement by HPLC

To measure DA levels in organoid lysates, high-performance liquid chromatography (HPLC) with electrochemical detection, was employed. Organoids were incubated in physiological medium containing 140 mM NaCl; 2.5 mM KCl; 1 mM MgCl2; 1.8 mM CaCl2 and 20 mM HEPES at pH 7.4. To measure the release of neurotransmitters under high KCl concentrations, organoids were incubated in high KCl medium (previous physiological medium with 56LJmMLJKCl, the concentration of NaCl was proportionally reduced to keep the same osmolarity). Solutions were collected after 20 minutes of incubation. The incubation solutions and the homogenates were centrifuged at 20,000 g for 15 minutes at 4°C. Supernatants were collected for analysis and were mixed with stabilization medium resulting in a final concentration of 250 µM ascorbic acid, 0.05 mM Na**_2_**EDTA and 4 mM formic acid. DA content was measured using an HPLC system equipped with an electrochemical detector (Agilent Technologies). Separation was achieved on a C18 reversed-phase column with a mobile phase consisting of 0.1 M phosphate buffer (pH 3.0), 0.1 mM EDTA, and 10% methanol at a flow rate of 1.0 ml/min. DA was quantified by comparing peak areas with those of standard solutions and normalized according to the protein quantification of individual organoid lysates.

### MEA analysis

Electrical activity was recorded from four midbrain organoids per condition at DIV 42 using the BioCAM DupleX system (3Brain, Pfaeffikon, Switzerland) with Accura-3D HD-MEA chips (4096 micro-needle electrodes, 90 µm height, 64 × 64 grid). The medium was changed 48 hours prior to recording. Organoids were placed on the electrodes in the Accura-3D recording chamber. After a 10-minute equilibration at 37°C in a miniature incubator, baseline signals were acquired for 10 minutes at a 20 kHz sampling rate (100 Hz high-pass filter). Organoids were then treated with 100 µM 4-aminopyridine (4-AP, Sigma-Aldrich), and recordings continued for 5 minutes. Tetrodotoxin (TTX) was applied at the end to exclude artifacts. Spikes were detected using the Precise Timing Spike Detection (PTSD) algorithm with a threshold of 9× the noise standard deviation, combined with AI validation (BrainWave 5 software, 3Brain).

### α-Synuclein preformed fibrils

αSyn-PFFs were provided by the Aligning Sciences Across Parkinson’s (ASAP) initiative through the laboratory of Laura Volpicelli-Daley and prepared according to the published guideline ^83^. For experiments employing fluorescently labelled αSyn-PFFs, fibrils were conjugated according to manufacturer’s instructions with the PE/Atto 594 Conjugation Kit – Lightning-Link^®^ (Abcam, ab269900). In an additional step, to remove residual unbound dye, αSyn-PFF 594 was centrifuged 3 times at 15000g for ten minutes each and washed with 1 mL of PBS after every centrifugation step. αSyn-PFF-594 were resuspended in 100uL of TRIS-Saline-Buffer in the final step. For both labelled and unlabelled, αSyn-PFFs were sonicated in the Qsonica700 sonicator (Qsonica L.L.C, CT, USA) equipped with a chilled cup horn at 15°C and handled according to an established protocol^84^. For the sonication, the following parameters were used: Sonication pulse 3 seconds on and 2 seconds off; 30% amplitude for 15 minutes followed by an additional sonication for 7.5 minutes. For all experiments, αSyn-PFFs were sonicated directly before treatment of the organoids.

### Treatment of midbrain organoids with **α**Syn-PFFs

Midbrain organoids were generated according to the tri-phasic WNT modulation protocol and bioreactor cultivation method described here. At DIV 80, organoids were selected for αSyn-PFF treatments. Individual WntTriP-Bio organoids were carefully transferred from the bioreactor using a cut pipette tip to a 48-well cell culture plate (Corning, NY, USA). To ensure homogeneous treatment conditions, one organoid was placed in one well each. For treatments, organoids received either αSyn-PFFs at a final concentration of 5μg/ml or an equivalent volume of TRIS-Saline-Buffer as controls. To ensure uniform exposure to αSyn-PFFs, organoids were gently flipped daily and maintained under static conditions for 48 hours at 37°C in a humidified incubator with 5% CO2. After 48 hours, organoids were transferred back to the bioreactor and supplemented with fresh medium. To account for any potential stress or damage during handling, TRIS-Saline-Buffer treated control organoids underwent identical transfer to 48-well-plates, medium changes, and manipulation procedures throughout the experiment. Organoid culture medium was refreshed every 3 days.

### Statistical Analysis

Data were analyzed using GraphPad Prism Software (Win10, version 10.2.2) or R. Statistical significance for imaging and bulk assays was determined by two-tailed t-test or one-way/two-way ANOVA, followed by Tukey’s post hoc test for multiple comparisons, as indicated. For single-cell analyses that incorporated biological replicate structure, linear mixed-effects models (LMM) were used for continuous outcomes, including score-based and pseudotime-derived comparisons, whereas generalized linear mixed-effects models (GLMM) were used for count- or class-based outcomes, including label-transfer analyses, with sample included as a random effect where appropriate. At least three organoids were analyzed per condition and time point. Differences were considered statistically significant at p < 0.05.

## Author contribution

H.R. and M.D. conceived and designed the experiments. H.R., F.B., and A.L. contributed to cell culture work. H.R., F.B. and M.J.P. performed and analyzed scRNA-seq. L.V.D. generated α - synuclein preformed fibrils and provided guidance on α-synuclein experiments. H.R. and M.D. wrote the manuscript with contributions from all authors. All authors read and approved of the final manuscript.

## Acknowledgments

We acknowledge the funding support of ERC CoG (#101003329, to M.D.), Fondation de France (# 00147852/WB-2023-51647, to M.D.) and Fondation pour la Recherche Médicale (FRM # MND202411019762, to M.D.). HR was supported by a UPC BiosPC scholarship (contrats doctoraux d’état #D25ZNWDC). Electrophysiological experiments were performed by the ePHYS core facility, led by Carine Dalle, at the Paris Brain Institute (ICM). This research was funded by Aligning Science Across Parkinson’s [Grant number: ASAP-000420] through the Michael J. Fox Foundation for Parkinson’s Research (MJFF) (to M.D.) and by additional funding support through MJFF (# 984115, to L.V.D.). An earlier version of this manuscript was posted to bioRxiv on July 30, 2025, at the following link: https://doi.org/10.1101/2025.07.29.667404. For the purpose of open access, the author has applied a CC BY 4.0 public copyright license to all author accepted manuscripts arising from this submission.

## Data availability

The data, code, protocols, and key lab materials used and generated in this study are listed in a Key Resources Table alongside their persistent identifiers at the following link: https://doi.org/10.5281/zenodo.20039098. Raw single-cell RNA-seq data are available in GEO under accession number GSE301365.

**Supplementary information is available on the Molecular Psychiatry website.**

