## Supplementary Figures for "Engineering functional ventral midbrain dopaminergic neurons in human organoids through WNT modulation and bioreactor culture"

### Supplementary figure legends

**Supplementary Figure 1. Generation and characterization of human midbrain organoids through tri-phasic modulation of WNT signaling.**

The effects of WntTriP-Bio culture conditions on dopaminergic maturation and astroglial representation were assessed by immunofluorescence across two additional, independent human control induced pluripotent stem cell lines (C2 and C3). **(A)** Brightfield images showing the morphology of human midbrain organoids (hMOs) at DIV0, DIV7, DIV8, DIV11, and DIV30. Representative organoids at each time point illustrate neuroepithelial bud formation and progressive growth. Scale bars: 500 µm. **(B)** Immunofluorescence images of DIV20 hMO cryosections cultured under Standard-OS or WntTriP-OS conditions. Sections were stained for FOXA2 (green) and TUBB3 (red), SOX2 (green) and MAP2 (red), OTX2 (green) and MAP2 (red), or TH (red). Nuclei were counterstained with DAPI. Scale bars: 50 µm.

**Supplementary Figure 2. Single-cell transcriptomic analysis reveals enhanced ventral midbrain and substantia nigra pars compacta-like dopaminergic specification following tri-phasic WNT modulation.**

**(A)** Label transfer of the Braun et al. human fetal reference onto DIV25 human midbrain organoids (hMO) cells, colored by predicted post-conception week (PCW), shows predominant alignment with early second-trimester stages. **(B)** Stacked bar plots of Seurat cluster proportions across individual hMOs at DIV25, DIV40, and DIV65, indicating consistent cellular composition across replicates and time points. **(C)** CellChat analysis of non-canonical WNT (*WNT5A*) and FGF signaling at DIV25 in Standard-OS and WntTriP-OS organoids reveals pathway reorganization in WntTriP conditions around a floor plate-centered *SOX6*$^{+}$/*EBF1*$^{+}$ ventral midbrain dopaminergic

(VM DA) niche. **(D)** Bubble plot of top differential ligand-receptor interactions involving the *SOX6*$^{+}$/*EBF1*$^{+}$ VM DA cluster in WntTriP versus Standard organoids, grouped by axon guidance, synaptic/adhesion, and VM patterning. Arrows denote interaction directionality relative to the *SOX6*$^{+}$/*EBF1*$^{+}$ cluster. **(E)** Pathway directionality analysis in WntTriP organoids highlighting shared incoming and outgoing signaling associated with the *SOX6*$^{+}$/*EBF1*$^{+}$ VM DA cluster.

**Supplementary Figure 3. Spinning bioreactor culture reduces apoptosis and enhances ion-channel-associated transcriptional programs in human midbrain organoids.**

**(A)** UMAP of single-cell transcriptomes from Standard-OS and Standard-Bio human midbrain organoids (hMOs) at DIV65, colored by cluster. **(B)** UMAP from Standard-OS and Standard-Bio hMOs at DIV65, colored by culture condition. **(C)** Feature plots showing expression of radial glia (*SOX2*), neuronal progenitor (*NHLH1*), and neuronal (*DCX*) markers in Standard-OS and Standard-Bio hMOs. **(D)** UCell apoptosis scores projected onto UMAP, visualized as cellular density. **(E)** Violin plots comparing apoptosis scores between Standard-OS and Standard-Bio hMOs at DIV65. **(F, G)** UCell ion-channel scores visualized on UMAP (F) and compared by violin plots (G) across conditions. **(H)** UCell synaptic and neuronal activity scores projected onto UMAP as cellular density.

**Supplementary Figure 4. WntTriP-Bio supports dopaminergic neuronal maturation without altering astroglial representation.**

**(A)** Representative immunofluorescence images of DIV40 hMO sections from control lines C2 and C3 cultured under Standard-OS and WntTriP-Bio conditions, stained for VMAT2 (green), TH (red), MAP2 (cyan), and DAPI (blue). Boxed regions are shown at higher magnification in adjacent panels. Scale bars: 250 µm (overview) and 25 µm (insets). **(B)** Representative immunofluorescence images of GFAP (red) and DAPI (blue) in DIV65 hMOs across Standard-OS, Standard-Bio, WntTriP-OS, and WntTriP-Bio conditions, showing comparable astroglial signal across conditions. Scale bars: 50 µm.

**Supplementary Figure 5. WntTriP-Bio enhances synaptic protein abundance and dopaminergic neuron maturation in human midbrain organoids.**

**(A)** Immunoblot analysis of βIII-Tubulin and Synaptophysin in individual DIV65 human midbrain organoids (hMOs) cultured under Standard-OS and WntTriP-Bio conditions. Densitometric quantification of Synaptophysin normalized to βIII-Tubulin is shown below. Mean ± SD; unpaired two-tailed t-test; *p = 0.0319. **(B)** Representative reconstructions of TH+ dopaminergic neurons from DIV100 organoid sections. White traces indicate reconstructed dopaminergic processes. Low-magnification TH images and corresponding TH/DAPI overlays are shown. Scale bars: 25 µm. **(C)** Quantification of filament and dendrite lengths (µm) of TH+ neurons in organoid sections under Standard-OS and WntTriP-Bio conditions. Unpaired two-tailed t-test, ****p < 0.0001. Maximum number of detected filaments + dendrites: Standard-OS, 498; WntTriP-Bio, 1284. Tile-stitched whole-organoid images were used; n = 3 organoids per condition. **(D)** Quantification of branching segments in TH+ neurons from organoid sections under Standard-OS and WntTriP-Bio conditions, showing increased arborization complexity in WntTriP-Bio DA neurons. Number of detected filaments analyzed: Standard-OS: 573; WntTriP-Bio: 608.

**Supplementary Figure 6. WntTriP-Bio organoids enrich an EBF1-high *SOX6+*/*KCNJ6+* substantia nigra-like dopaminergic program associated with synaptic maturation and reduced cellular stress.**

**(A)** Heatmap showing per-sample neuronal cluster composition in DIV65 Std-OS and WntTriP-Bio human midbrain organoids (hMOs), highlighting reproducible clustering across biological replicates. Values indicate the percentage of cells assigned to each cluster per organoid. **(B)** Feature plots showing expression of *SOX6*, *LMX1B*, *EBF1*, *SV2C, CALB1, and FOXP2* in the DIV65 neuronal subset, illustrating enrichment of SNpc-associated neuronal features in WntTriP-Bio hMOs, and VTA-associated enrichment in Standard-OS. **(C)** Dot plot summarizing the relative expression of representative VTA-like and SNpc-associated markers across conditions in the

DIV65 neuronal subset. **(D)** Per-sample heatmap of pathway-associated genes across Std-OS and WntTriP-Bio organoids. Genes linked to synapses, ion channels, and axon/projection programs are increased in WntTriP-Bio organoids, whereas stress- and apoptosis-associated genes are relatively enriched in Std-OS hMOs. **(E)** Violin plots of module scores for synapse, stress and apoptosis, and neurotransmitter/ion-channel gene sets across conditions. **(F)** PySCENIC regulon analysis showing differential regulon activity between Std-OS and WntTriP-Bio organoids. The left panel summarizes regulon shifts, and the right panel shows the activity distribution of the EBF1 regulon, which is increased in WntTriP-Bio organoids. **(G)** Gene Ontology biological process enrichment analysis in the *EBF1*+/*SOX6*+/*KCNJ6*+ DA cluster 1, highlighting pathways related to axon development, synapse organization, trans-synaptic signaling, cell projection, neuron migration, and neuron cell-cell adhesion.

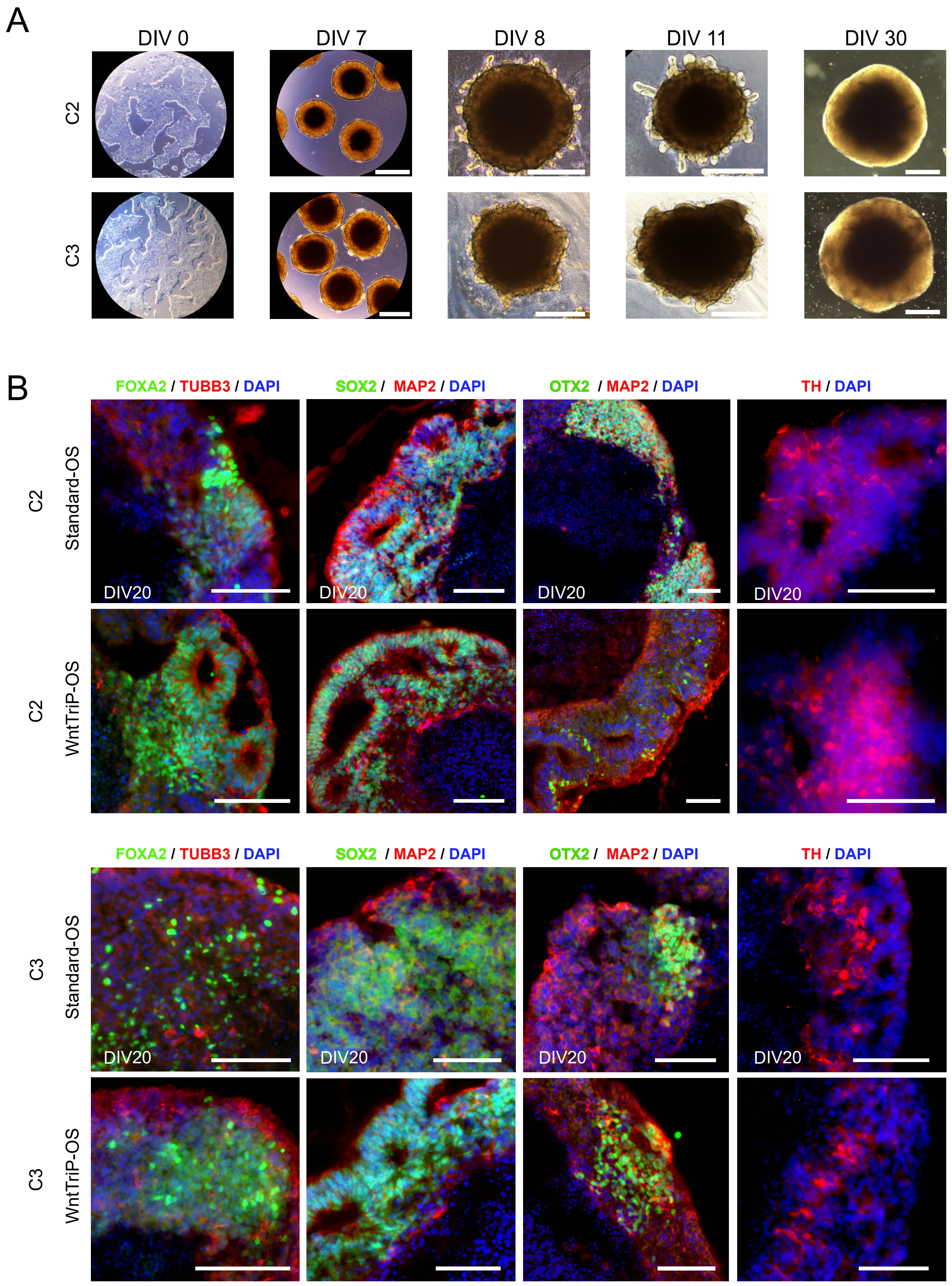


Supplementary Figure 1

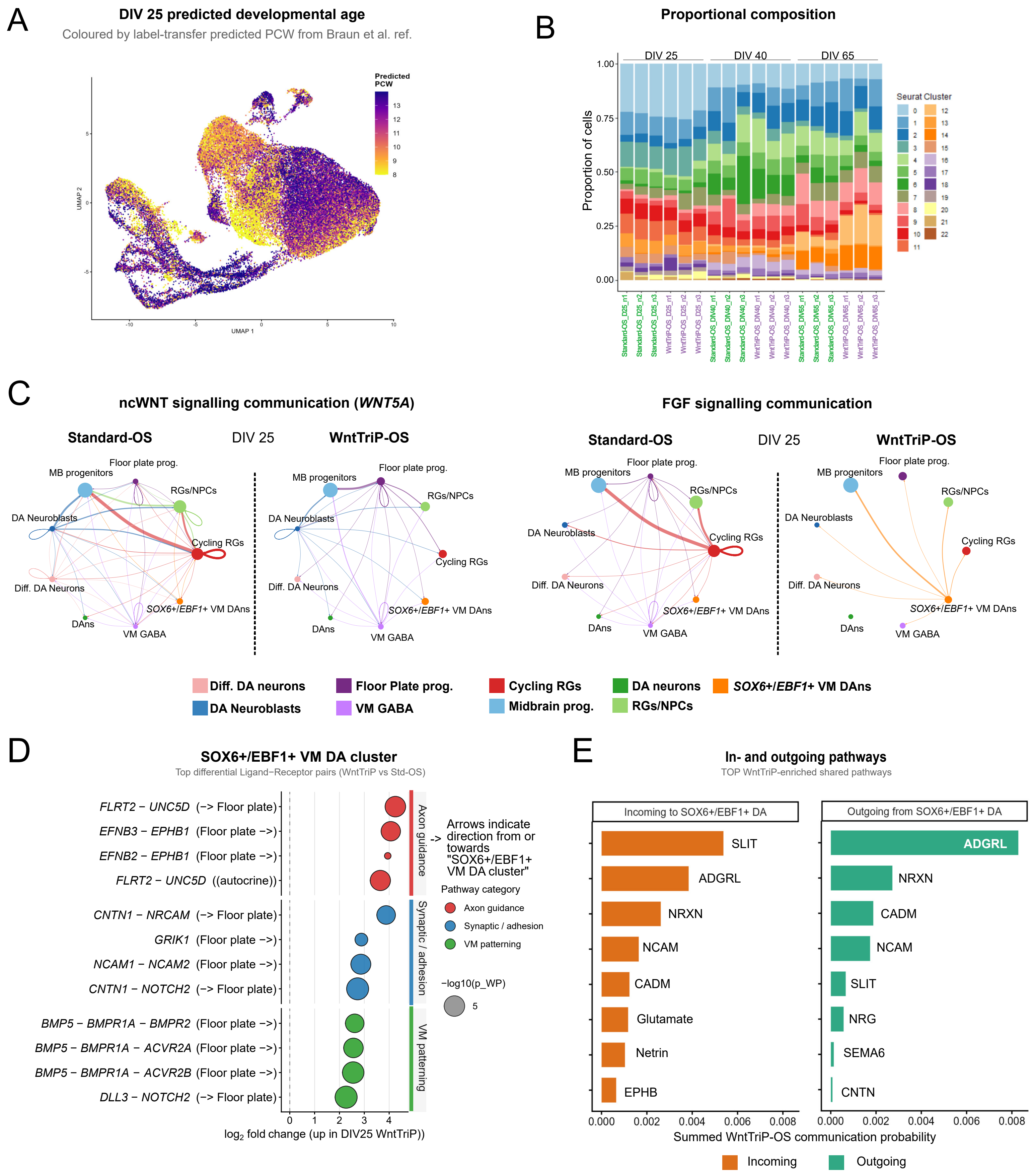


**Supplementary Figure 2**

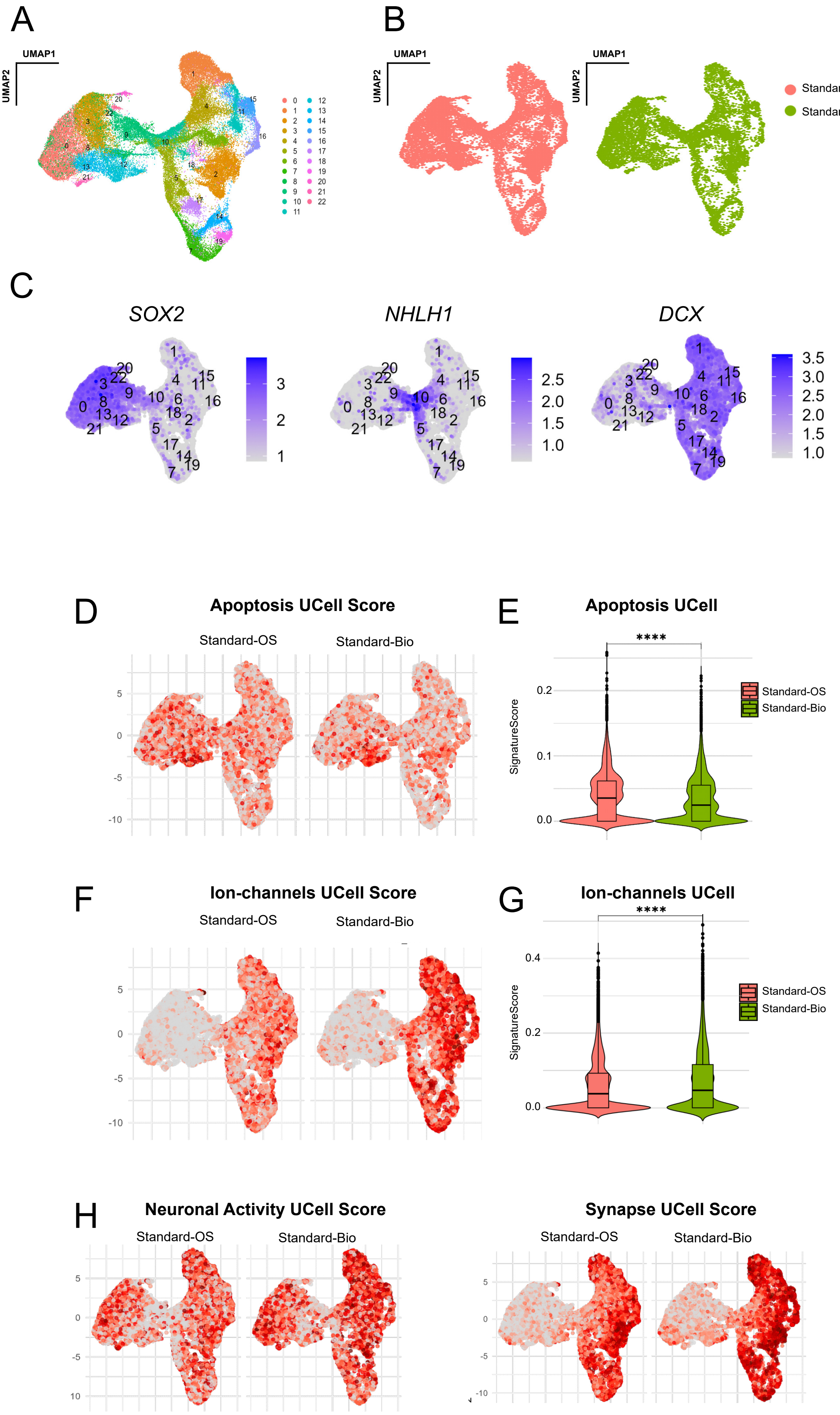


**Supplementary Figure 3**

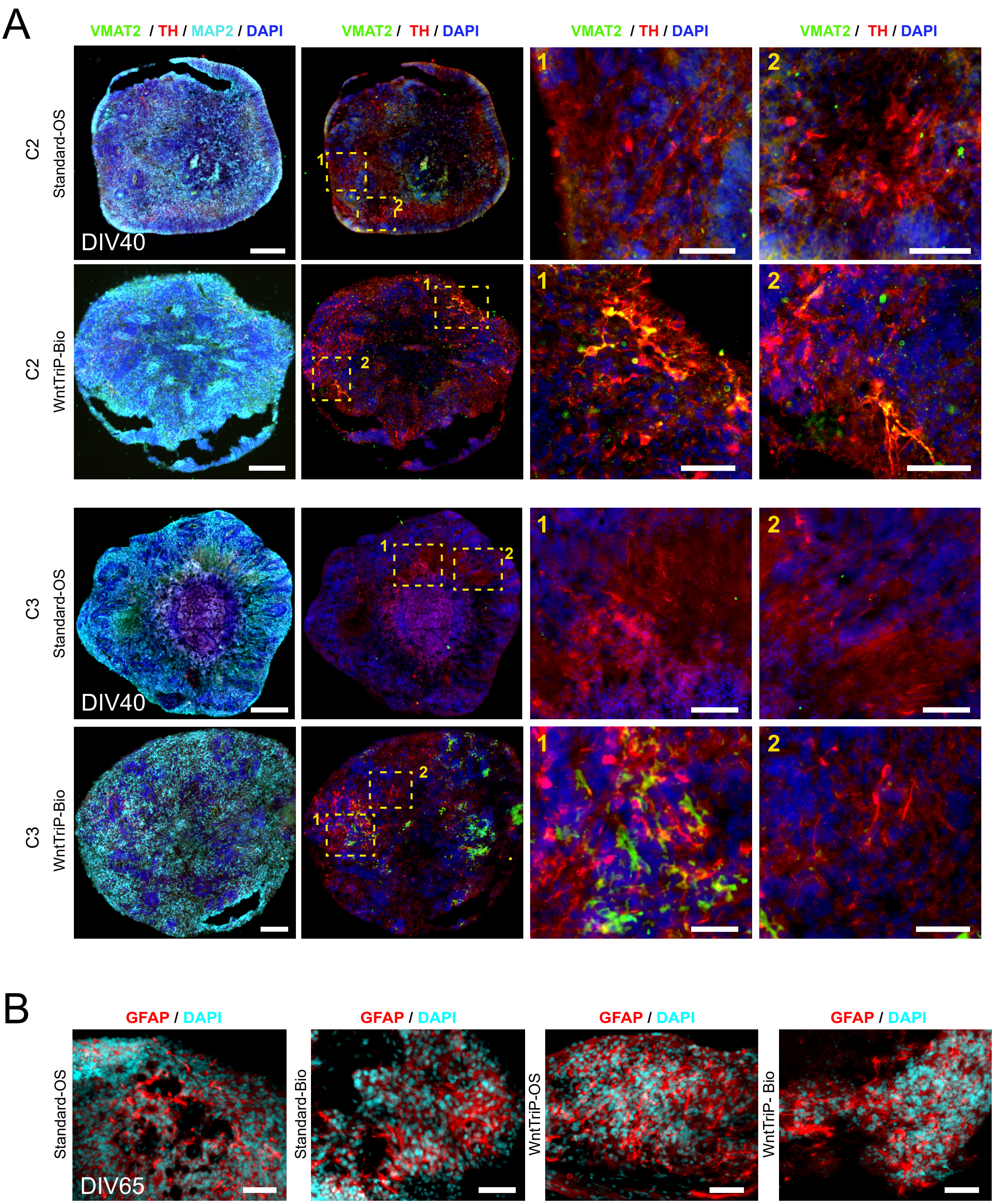


**Supplementary Figure 4**

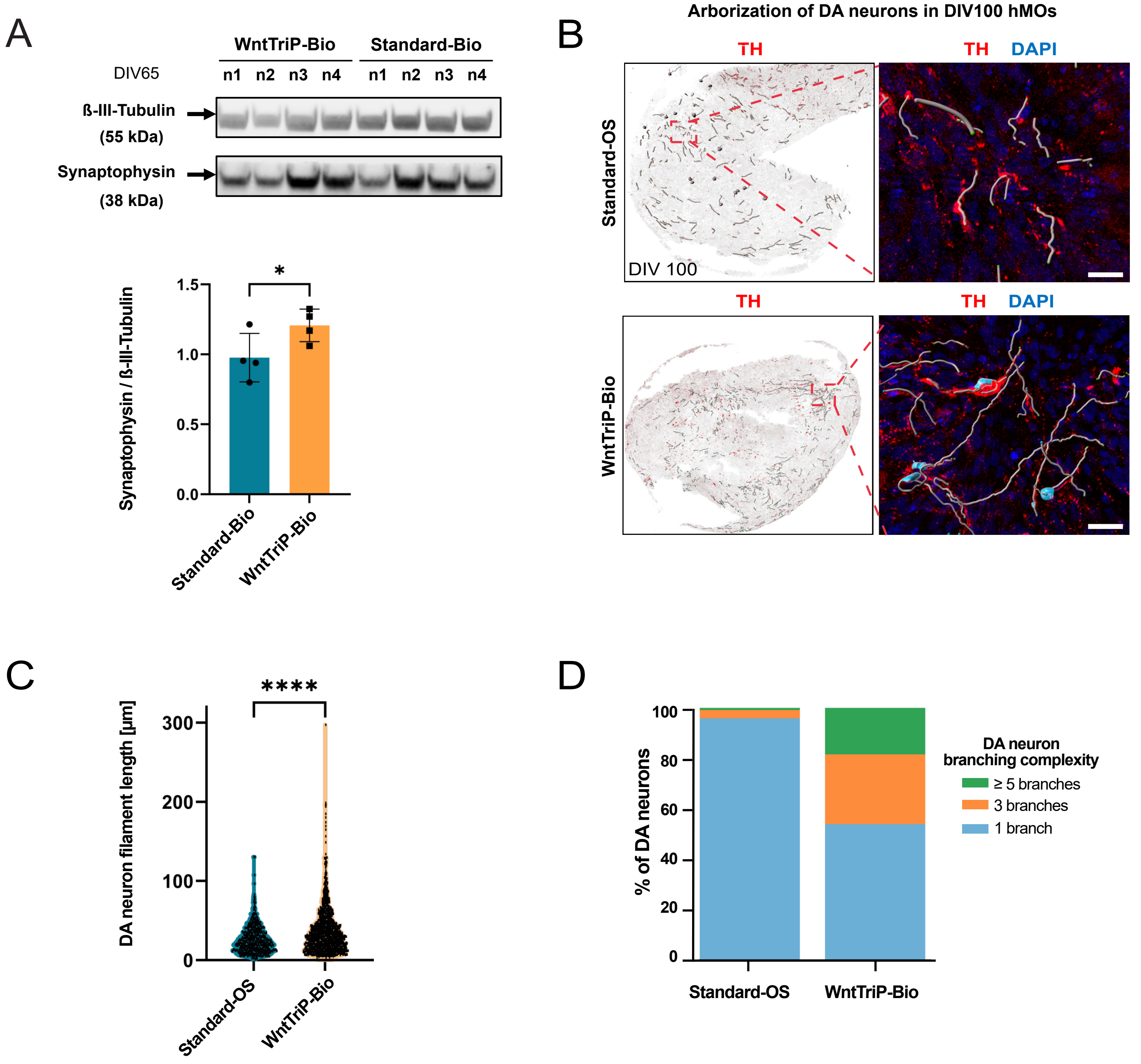


**Supplementary Figure 5**

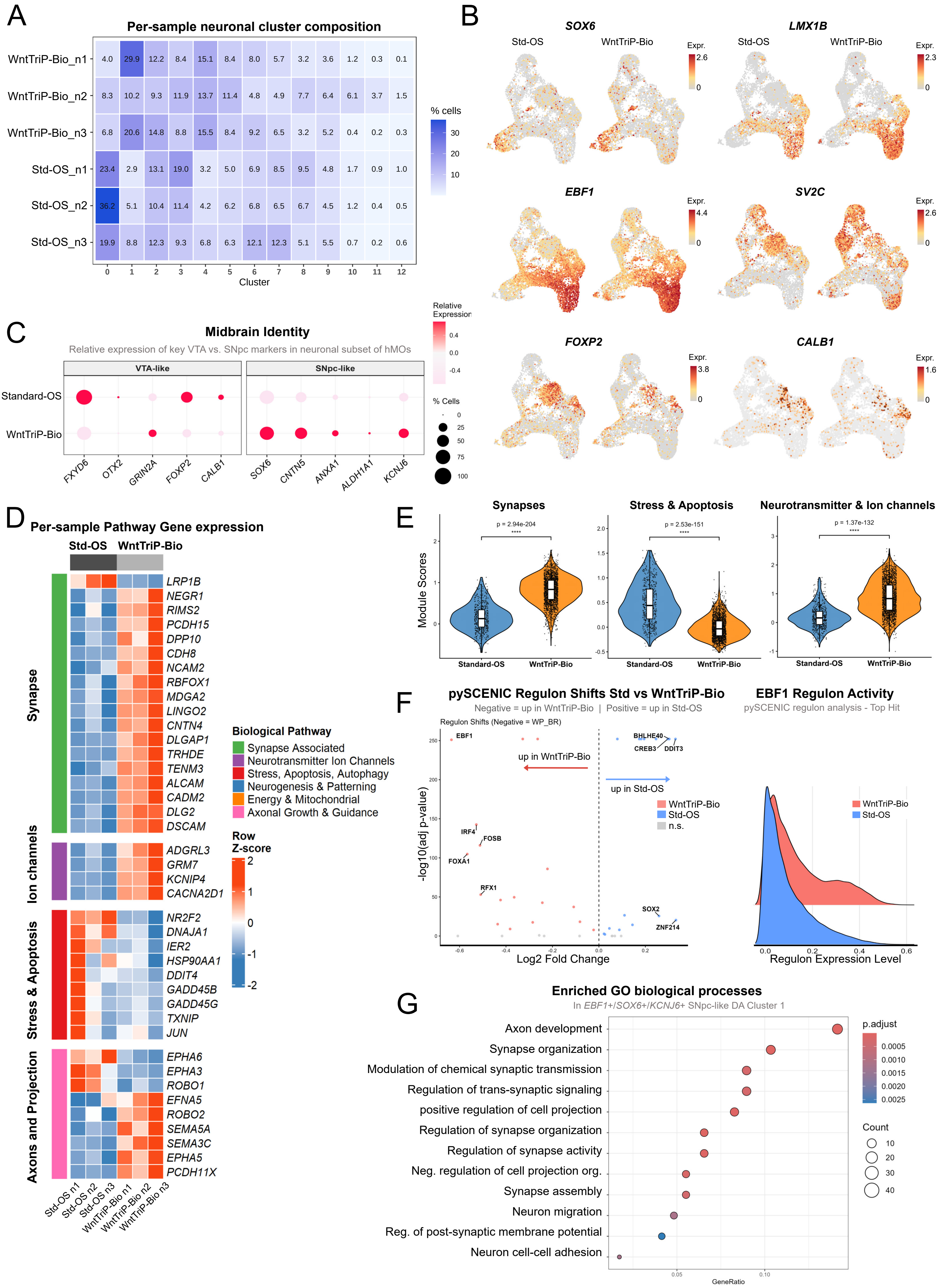


**Supplementary Figure 6**
